# Lipid nanoparticle shape and morphology determine the magnitude and duration of mRNA transfection

**DOI:** 10.64898/2026.10.18.759824

**Authors:** Sara Elsafy, Martin Dulle, Saba Shahzad, Thomas Hammoor, Marek Weiler, Ben Peacock, Joseph Brealey, Josbert M. Metselaar, Twan Lammers, Alexandros Marios Sofias

## Abstract

Lipid nanoparticles (LNP) are widely used for nucleic acid delivery, with current optimization efforts mainly focusing on refining formulation features, enabling extrahepatic targeting, and improving therapeutic efficacy. Particle shape and internal architecture are variables often overlooked in LNP design, although they critically influence particle stability, biodistribution, cellular uptake, and transfection. Here, we investigate how formulation and process parameters impact LNP shape, morphology, and internal structure, and how these structural features, in turn, influence mRNA loading, cell uptake, and expression. We demonstrate that modulation of lipid composition by increasing DSPC content drives a transition from compact spherical to vesicular non-spherical morphologies, with altered RNA loading at the single-particle level. While non-spherical LNP exhibited higher cellular uptake than spherical LNP, it was identified that this increase was relatively modest compared with the substantially higher and sustained protein expression for non-spherical structures. This suggests that increased uptake alone cannot explain the enhanced transfection performance. The trend was conserved across LNP formulations comprising clinically relevant ionizable lipids SM-102, ALC-0315 and DLin-MC3-DMA. Taken together, these findings establish LNP shape and internal morphology as functionally relevant design variables that should be considered in the design of next-generation LNP with improved mRNA delivery and expression.

**Graphical abstract:** 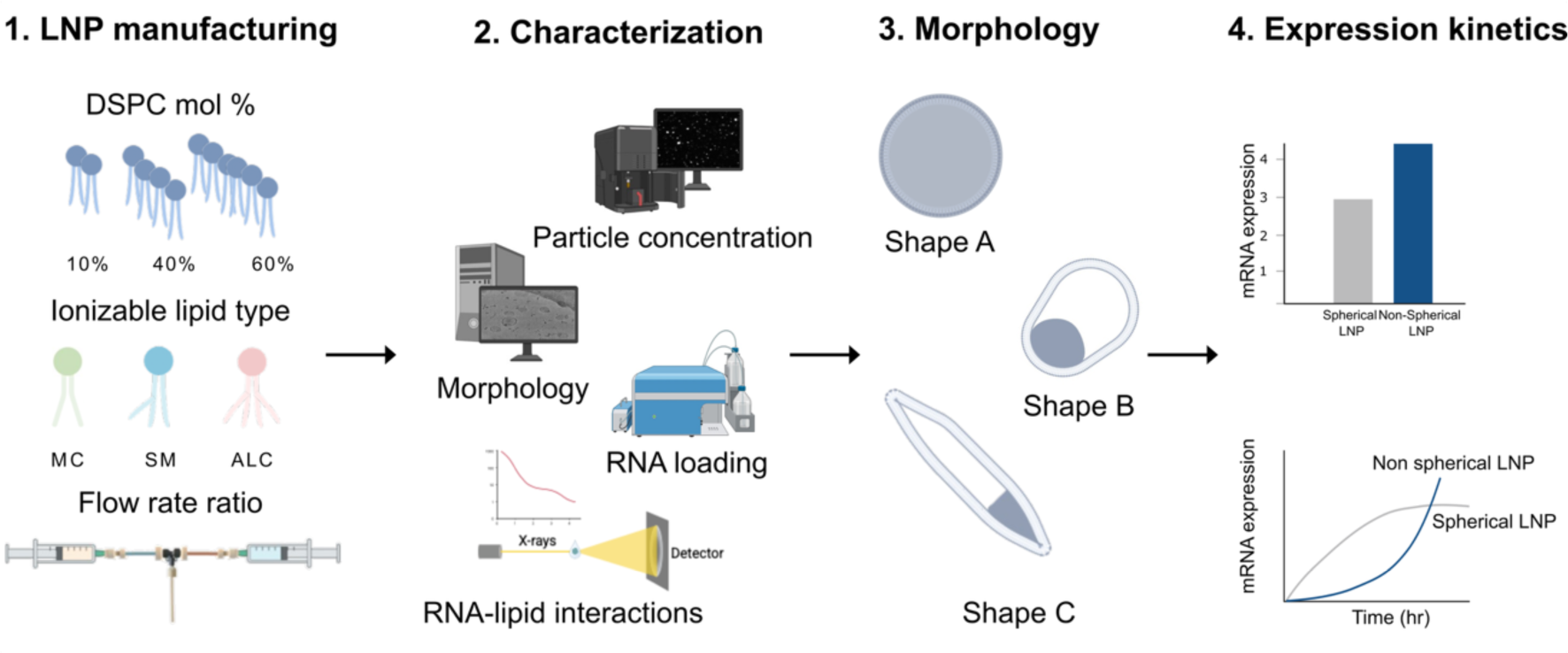

## 1. Introduction

Lipid nanoparticles (LNP) have become the leading non-viral vectors for nucleic acid delivery^1^. LNP protect nucleic acids from degradation, enhance their cellular uptake, and enable precise gene modulation within target cells^2^. Their clinical success was first demonstrated by the FDA approval of Patisiran (Onpattro^™^), the first siRNA-LNP to treat hereditary transthyretin-mediated (hATTR) amyloidosis, a rare genetic disease characterized by the accumulation of misfolded transthyretin (TTR) protein deposits in various organs and tissues^3^. This breakthrough was followed by the approval of two COVID-19 mRNA LNP vaccines (Moderna’s Spikevax^™^ and Pfizer/BioNTech’s Comirnaty^™^; which encode the SARS-CoV-2 spike protein), and more recently the respiratory syncytial virus (RSV) vaccine (Moderna’s mRESVIA^®^; encoding prefusion F glycoprotein)^4–6^. Building on this success, LNP are being actively explored for applications beyond vaccines, including cancer therapies, protein replacement, and gene-editing^7–9^.

LNP are composed of four key lipid components: (i) an ionizable cationic lipid for facilitating RNA encapsulation and promoting endosomal escape, (ii) a phospholipid for supporting and stabilizing the LNP structure, (iii) cholesterol for enhancing particle stability and improving cellular uptake, and (iv) a PEG-lipid for controlling LNP size, preventing aggregation, and improving circulation half-life^2^. These components play a crucial role in determining stability, RNA loading, in vivo biodistribution, cellular uptake, and finally transfection efficiency. As such, lipid composition is a critical factor to consider during the design and manufacturing of LNP. In addition to formulation composition, microfluidic manufacturing parameters, such as flow rate ratio (FRR) and total flow rate (TFR), are well recognized for influencing the physicochemical properties of LNP^10–14^.

Given the central role of lipid composition in determining LNP performance, substantial efforts have focused on optimizing the chemical structure of ionizable lipids to enhance potency and enable extrahepatic delivery^15–18^. In parallel, modifications to helper lipids (cholesterol and phospholipids) have been explored as complementary strategies to improve LNP efficacy and (extrahepatic) tissue targetability^19–22^. These studies primarily aimed to elucidate the contribution of individual lipid components to the biological efficacy of LNP formulations. To isolate these effects, lipid molar ratios comparable to those in Onpattro^™^ were often used and fixed across formulations, yielding LNP with comparable particle sizes and morphology.

Nevertheless, emerging evidence suggests that major changes in LNP performance often arise from pronounced shifts in particle morphology, driven either by the incorporation of structural lipids and cholesterol analogs or by substantial alterations in the molar ratios of phospholipids in the LNP formulation^19–23^. Such structural deviations have been associated with enhanced mRNA delivery^22,24^ or modified accumulation in extrahepatic tissues^21,22,24–26^. These observations are consistent with the broader understanding that nanoparticle shape can strongly influence particle stability, protein corona formation, cellular uptake mechanisms, intracellular processing, and ultimately therapeutic outcome^17,22–24,26–28^.

Based on these findings, it is evident that particle shape and morphology are largely treated as a secondary consequence of formulation conditions rather than as an independent, tunable design variable. They are, thus, rarely examined for their functional implications. As a result, the role of LNP shape in governing RNA loading, cellular uptake, intracellular trafficking, and mRNA expression kinetics remains poorly defined.

In this study, we aimed to systematically investigate how formulation and process parameters govern LNP shape and how these structural features, in turn, influence mRNA loading, delivery, and expression. Using a combination of biophysical characterization and functional assays, we demonstrated that non-spherical LNP exhibit distinct mRNA expression kinetics characterized by enhanced and sustained protein expression despite reduced RNA loading.

## 2. Materials and methods

### 2.1. Materials

DSPC 18:0 PC (1,2-distearoyl-sn-glycero-3-phosphocholine) and DMG-PEG(2K) (1,2-dimyristoyl-rac-glycero-3-methoxypolyethylene glycol-2000) were obtained from Avanti^®^ Polar Lipids (Merck KGaA, Germany). Ionizable lipids SM-102, ALC-0315 and DLin-MC3-DMA were purchased from MedChemExpress (Hycultec GmbH, Germany). Cholesterol, sodium acetate, and poly A were purchased from Sigma-Aldrich (Merck KGaA, Germany). DiR DiIC18(7) (1,1’-Dioctadecyl-3,3,3’,3’- Tetramethylindotricarbocyanine Iodide) dye, Lipofectamine^™^ MessengerMAX^™^ transfection reagent, and Quant-it^™^ RiboGreen RNA Assay Kit were acquired from Invitrogen^™^ (Thermo Fisher Scientific, Germany). Spectra/Por^®^ regenerated cellulose dialysis membrane tubing (MWCO 12-14 kDa) was obtained from Spectrum™ Labs (Thermo Fisher Scientific, Germany). Phosphate-buffered saline (PBS) ready-to-use tablets (ROTI^®^Fair PBS 7.4) were purchased from Carl Roth^®^ (Carl Roth GmbH, Germany). Amicon^®^ Ultra Centrifugal filter (10 kDa MWCO) and 0.2 μm disposable polyethersulfone Nalgene^®^ syringe filter units were obtained from Merck Millipore (Merck KGaA, Germany). Cholesterol FS Assay and cholesterol standard were obtained from DiaSys (Diagnostic Systems GmbH, Germany). HiScribe T7 mRNA Kit with CleanCap^®^ Reagent AG and Monarch^®^ RNA Cleanup Kit were obtained from New England Biolabs (New England Biolabs GmbH, Germany). Cell culture medium and supplements were obtained from Gibco (Thermo Fisher Scientific, Germany). All other reagents were of analytical grade. PHD ULTRA^™^ advanced programmable syringe pumps were purchased from Harvard Apparatus (Holliston, USA).

### 2.2. Design of Experiments (DoE)

A custom experimental design was employed to systematically explore the influence of formulation parameters on LNP physicochemical properties. Lipid composition (DSPC mol%), FRR, and lipid-to-nucleic acid ratio (N:P ratio) were selected as key variables and varied within predefined ranges. Specifically, DSPC ranged from 10 to 70 mol %, FRRs were set at 3, 4, and 5, and N:P ratios were set to 3, 6, or 12. A constrained number of 10 runs was selected to efficiently sample the multidimensional parameter space while minimizing the number of experimental runs. The detailed list of experimental conditions is provided in **Table S1.** Each run was prepared, and the LNP size, polydispersity index (PDI), and particle concentration were measured by dynamic light scattering [DLS; Zetasizer Nanoseries ZS90 (Malvern Instruments Ltd., UK)] and nanoparticle tracking analysis [NTA; NanoSight system (Malvern Instruments Ltd., UK)]. Experimental data were analyzed using JMP software (SAS Institute, USA), and response surface plots were generated to visualize trends in particle size, concentration, and dispersity as a function of formulation parameters. Given the limited number of experimental runs, the design was intended to identify general trends rather than to construct a fully predictive statistical model.

### 2.3. LNP Production

#### 2.3.1. In vitro transcription (IVT) of enhanced green fluorescent protein (eGFP) mRNA

An eGFP coding sequence with a 30 + 70-nucleotide poly A tail was cloned into a plasmid, amplified, sequenced, and linearized with HindIII. The linearized template was transcribed using the HiScribe CleanCap^®^ IVT kit (New England Biolabs GmbH, Germany), with CleanCap AG trinucleotide cap 1 analog incorporated as a co-transcriptional cap structure. All uridine residues in the transcript were fully substituted with N1-methylpseudouridine (m1Ψ) for improved stability and reduced immunogenicity. Prior to starting, all kit components were thawed on ice and briefly centrifuged at 2,000 g for 30 seconds. Reagents were added to an RNase-free Eppendorf tube in the order and volumes specified in the manufacturer’s manual. The reaction mixture was then incubated at 37 °C for 6 hours. Following transcription, the IVT-mRNA was purified using the Monarch^®^ RNA Cleanup Kit (New England Biolabs GmbH, Germany) according to the manufacturer’s instructions. The integrity of the purified mRNA was assessed via gel electrophoresis, and the yield was quantified using a NanoDrop One^C^ spectrophotometer (Thermo Fisher Scientific, Germany).

#### 2.3.2. Preparation of RNA LNP

RNA LNP were produced using a previously described method^29^. Briefly, the organic phase was prepared by dissolving ionizable lipid, DSPC, cholesterol, and DMG-PEG2000 at an initial lipid concentration of 12 mM. The aqueous phase consisted of either eGFP mRNA or poly A dissolved in 25 mM sodium acetate buffer (pH 4) at an N:P ratio of 6. The organic and aqueous solutions were loaded into syringes, placed on a PHD ULTRA^™^ programmable syringe pump (Harvard Apparatus, USA), and passed through a T-junction microfluidic mixer^12,29,30^. The FRR and TFR were fixed at 3:1 and 20 mL/min, respectively. The specific lipid molar ratios, N:P ratios, TFR, and FRR of LNP prepared for the DoE are summarized in **Tables S1 and S2**. For cell transfection assays, DiR-labelled LNP were prepared by incorporating 0.2 mol% DiR (Invitrogen^™^, Thermo Fisher Scientific, Germany) into the formulation. To maintain a constant total lipid molar percentage, the addition of the DiR fraction was compensated by a corresponding reduction in cholesterol content. Following formulation, LNP were dialyzed against PBS (pH 7.4) using Spectra/Por^®^ regenerated cellulose dialysis membrane tubing (MWCO 12-14 kDa) with stirring at 200 rpm for 24 hours to remove residual ethanol. After dialysis, the formulation was sterile filtered using a 0.2 μm Nalgene^®^ syringe filter, transferred to an Amicon^®^ filter (MWCO 10 kDa), and centrifuged at 3000 x g for 40 minutes (or until the volume was concentrated to 250 μl from an initial 2 ml). The concentrated LNP were stored at 4°C until further use. All experiments were conducted using RNase-free consumables and buffers.

### 2.4. LNP characterization

#### 2.4.1. Hydrodynamic diameter (Z-average) and Polydispersity Index (PDI)

LNP size (Z-avg) and PDI were measured using DLS on a Zetasizer Nano ZS90 (Malvern Instruments Ltd., UK). Concentrated LNP samples were diluted 100-fold in PBS, and DLS measurements were conducted at 25°C after an initial equilibration period of 120 seconds. Each sample underwent three measurements, with the number of runs set automatically by the instrument. For in vitro stability assessment under physiologically relevant conditions, LNP were incubated in either PBS or a bovine serum albumin/PBS (BSA/PBS; 4.5% w/v BSA) solution at 37°C with shaking. DLS measurements were taken at 0, 1, 4, 8, and 24 hours to assess changes in particle size and PDI over time.

#### 2.4.2. Nanoparticle tracking analysis

Particle concentration was determined by NTA using a NanoSight NS300 system (Malvern Instruments Ltd., UK). RNA LNP samples were diluted at least 1:1000 in Milli-Q water prior to analysis and gently mixed to ensure homogeneity. Measurements were performed in light-scattering mode using a laser-based optical detection module (533 nm). For each sample, three sequential video acquisitions of 60 s duration were recorded and analyzed using NTA software (version 3.2 Dev Build 3.2.16). The camera level was set to 14 and the detection threshold to 7, while all other parameters were set to automatic. Identical acquisition and analysis parameters were applied across all samples to enable quantitative comparison of particle concentrations.

#### 2.4.3. Lipid content quantification

Cholesterol content was quantified using the DiaSys Cholesterol FS assay (Diagnostic Systems GmbH, Germany), an enzymatic colorimetric assay. A serial dilution of cholesterol standard (2 to 0.0156 mg/ml) was prepared in PBS, in triplicate, and placed in a transparent 96-well plate. Each LNP formulation was diluted 10x in PBS, and 10 μl of the diluted sample was added in triplicate to the 96-well plate. Next, 190 μl of the cholesterol reagent was added to both the standard curve wells and the LNP sample wells. The 96-well plate was incubated for 5 minutes at 37°C with shaking at 500 rpm. Absorbance was measured at 600 nm using a TECAN Infinite M200 PRO microplate reader (Tecan Trading AG, Switzerland), and cholesterol concentration in LNP was determined from the standard calibration curve. Total lipid concentration (mg/ml) was calculated using the following equation:

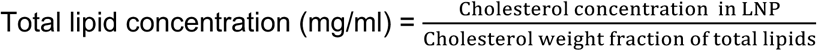

#### 2.4.4. RNA encapsulation

The amount of eGFP mRNA encapsulated in the LNP was quantified using the Quant-iT^™^ RiboGreen RNA Assay Kit (Thermo Fisher Scientific, Germany). A serial dilution of nucleic acid standards (10 to 0.078 µg/ml) was prepared in PBS, in triplicate, and added to a black 96-well plate. Concentrated LNP formulations were diluted 500-fold in PBS, and 50 μl of each diluted sample was added to six wells to measure both free and total mRNA in triplicates. Six additional wells containing 50 μl of PBS served as blanks. Two Tris-EDTA (TE) buffer solutions were prepared according to the manufacturer’s protocol: one with 2% (v/v) Triton reagent (TE + Triton) and one without. The RiboGreen fluorescent probe was then added to both buffers at a 1:100 dilution. Subsequently, 50 μl of the probe in Triton + TE buffer was added to the standard curve wells, three PBS blanks, and three LNP-containing wells. The remaining wells received 50 μl of the probe in TE buffer. The 96-well plate was incubated for 30 seconds with continuous shaking at 300 rpm. Fluorescence intensity (λ_Ex_ = 480 nm, λ_Em_ = 525 nm) was measured using a TECAN Infinite M200 PRO microplate reader (Tecan Trading AG, Switzerland). To account for background signal from the buffer and Triton, the fluorescence intensity from PBS wells with TE + Triton was subtracted from the fluorescence values of both the standard curve and LNP samples treated with TE + Triton, providing the signal from total RNA. Similarly, the fluorescence intensity from PBS wells containing TE buffer was subtracted from the fluorescence values of LNP samples treated only with TE buffer, yielding the signal from free (unencapsulated) RNA. The averages of the technical repeats were calculated, and the encapsulation efficiency percentage (EE%) was determined using the following equation:

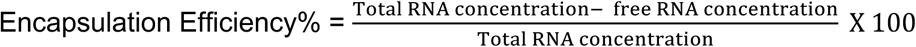

### 2.5. Cryogenic transmission electron microscopy (Cryo-TEM)

LNP morphology was assessed using CryoTEM. Lipid content of concentrated RNA LNP was determined using the DiaSys Cholesterol FS assay as described in Section 2.4.3. RNA LNP lipid concentration was adjusted to ∼ 8mg/ml prior to analysis. Sample vitrification was conducted using VitroJet system (DEMCON, formerly CryoSol-World B.V., The Netherlands). Initially, Quantifoil 1.2/1.3 Cu 200 grids with a 2 nm continuous carbon film (Quantifoil Micro Tools GmbH, Germany) were clipped into autogrid rings (Thermo Fisher Scientific, USA) at room temperature. The pre-clipped grids were loaded into the VitroJet system and plasma-cleaned for 60 s using the integrated plasma cleaner. Subsequently, 0.2 ul of LNP samples were deposited onto the grids using a pin-printing technique. The samples were vitrified by jetting liquid ethane onto the grids and subsequently transferred to and stored in liquid nitrogen at approximately −180 °C until imaging. Imaging was performed at the Ernst Ruska-Centre for Microscopy and Spectroscopy with Electrons (ER-C), Forschungszentrum Jülich GmbH, Jülich, Germany, using a Talos Arctica transmission electron microscope (Thermo Fisher Scientific, USA) operated at 200 kV and equipped with a Gatan K3 direct electron detector and Bioquantum energy filter. Images were acquired as movies using an automated EPU software at a nominal magnification of 100,000 x, corresponding to a calibrated pixel size of 0.8 Å/pixel with a total electron dose of 30 e⁻/Å². A nominal defocus of −5 µm was used, and the BioQuantum energy filter was operated with a 20-eV energy-selecting slit. Movie frames were gain-corrected and aligned using the Patch Motion Correction procedure in CryoSPARC^31^, the resulting motion-corrected micrographs were 2 x binned during preprocessing, yielding a final pixel size of 1.6 Å/pixel for subsequent image analysis. Scale bar was added using ImageJ software (NIH, USA)^13^. Following addition of the scale bar, ImageJ was used to measure the length and height of individual LNP based on the outer membrane boundaries. Particle length was defined as the longest distance from one side of the particle to the other. Particle width was measured perpendicular to the length, through the center of the particle, as the distance between the outer membrane boundaries. The aspect ratio of each particle was calculated by dividing its measured length by its width. Then the mean of the aspect ratios of all particles within each image was calculated. Three randomly selected cryo-TEM images were analyzed for each formulation (10-30 individual particles analyzed per image, yielding 30-90 particles in total per formulation). The mean aspect ratio from each image was plotted as a separate data point in GraphPad Prism (GraphPad Software, USA). For the quantification of blebbed particles, blebbed particles were defined as LNP displaying membrane protrusions surrounding an aqueous core in cryo-TEM images. For each image, individual LNP were classified as either blebbed or non-blebbed, and the percentage of particles displaying membrane blebs was calculated relative to the total number of particles analyzed. The same cryo-TEM images used for the aspect-ratio analysis were also used to quantify blebbed particles. The percentage of blebbed particles was determined for each image, and the average percentage across the three images was plotted using GraphPad Prism (GraphPad Software, USA).

### 2.6. Small Angle X-Ray Scattering (SAXS)

RNA LNP samples were prepared as described in Section 2.3, and scattering patterns were recorded using a “Ganesha-Air” SAXS system (SAXSLAB, Xenocs, USA). The X-ray source of this laboratory-based system was a D2-MetalJet (Excillum, USA) with a liquid-metal anode operating at 70 kV and 3.57 mA with Ga-Kα radiation (λ = 0.1341 nm) providing a very brilliant and a very small beam (<100 μm). The beam was focused with a focal length of 55 cm using a specially made X-ray optic (Xenocs, USA) to provide a very small and intense beam at the sample position. Measurements were done in 2.1 mm borosilicate glass capillaries (Hilgenberg GmbH, Germany) at RT. The scattering data were recorded by a position-sensitive detector (Eiger 4M, Dectris, USA) without a beam stop, covering the scattering vector range from 0.1 to 5 nm^−1^. The circularly averaged data were normalized to incident beam, counts per solid angle, and measurement time before subtraction of the appropriate buffer, PBS in this case, as background.

### 2.7. Nano flow cytometry (NanoFCM)

A NanoAnalyzer U30 instrument (NanoFCM Inc.) equipped with dual 488/638 nm lasers was used for simultaneous detection of side scatter (SSC) and fluorescence of individual particles. Single-photon counting avalanche photodiode detectors (SPCM APDs) with bandpass filters allowed for collection of light in specific channels (SSC – 488/10 nm; FL1 – 525/40 nm; FL2 – 670/30 nm). Gravity-fed HPLC-grade water served as the sheath fluid. Sampling pressure by air pump module was 1 kPa and measurements were taken over 1 minute. All samples were diluted in RNase-free 10 mM HEPES buffer, pH 7.4 to attain a particle count within the optimal range of 2,000 – 12,000/min. To assess RNA loading, fluorescent labelling of LNP with SYTO 9 membrane-permeable nucleic acid dye (Invitrogen, S34854) was performed. LNP were diluted to 2 x10^10^ particles/ml in HEPES and 9 μl was mixed with 1 μl SYTO 9 diluted to 100 μM in HEPES. Samples were incubated for 30 minutes at room temperature. Following incubation, samples were diluted in HEPES buffer to a final concentration of 1 x 10^8^ – 1 x 10^9^ particles/ml and analyzed immediately using the NanoAnalyzer. RNA quantification was performed following export of the raw data from NanoFCM Profession in CSV format. Then, LNP copy number was calculated as the fluorescence intensity of each mRNA-loaded LNP (SSC^+^ SYTO 9^+^) relative to the median fluorescence intensity of the free mRNA population (SSC^-^ SYTO 9^+^).

### 2.8. In vitro cell uptake and protein expression of eGFP mRNA LNP

#### 2.8.1. Flow cytometry

Human embryonic kidney 293T (HEK293T) and human cervical cancer (HeLa) cell lines were cultured in Dulbecco’s Modified Eagle Medium (DMEM; Gibco, Thermo Fisher Scientific, Germany). The medium was supplemented with 10% fetal bovine serum (FBS; Gibco, Thermo Fisher Scientific, Germany) and 100 U/ml penicillin–streptomycin (Gibco, Thermo Fisher Scientific, Germany). Cells were maintained at 37 °C in a humidified incubator with 5% CO₂. LNP samples were prepared as described in Section 2.3. For fluorescence tracking, 0.2 mol% of the lipophilic tracer DiR was added to the organic phase prior to mixing with mRNA. To evaluate cellular uptake and eGFP expression, 70,000 HEK293T cells and 50,000 HeLa cells were seeded in 24-well plates (Falcon/Corning, Corning, NY). After 24 hours, LNP were diluted in PBS to a concentration of 20 µg/ml, then added to 500 µl of fresh medium, resulting in a final dose of 200 ng mRNA per well. Cells were then incubated with the LNP samples for 24 hours. The tested groups included LNP formulated with varying DSPC and ionizable lipid compositions: MC-based LNP: 10%, 40%, and 60% DSPC, SM-based LNP: 10%, 40%, and 60% DSPC, ALC-based LNP: 10%, 40%, and 60% DSPC. Lipofectamine^™^ MessengerMAX^™^-treated cells served as a positive control, while PBS-treated cells served as a negative control. Following incubation, the medium was aspirated, and cells were washed with PBS. Afterward, cells were detached by incubation with 200 μl trypsin for 5 minutes at 37 °C, followed by deactivation of trypsin by adding four times the volume of complete medium. The cell suspension was transferred to FACS tubes, centrifuged at 400 g for 5 minutes, and the supernatant was removed. The cell pellet was washed twice, then resuspended in 500 µl of FACS buffer (2% FBS and 2 mM EDTA in PBS). LNP uptake and eGFP expression were analyzed using a BD LSRFortessa^™^ flow cytometer (BD Biosciences, USA), and BD FACSDiva^™^ Software. Data acquisition was performed for 10,000 gated viable events per sample, and analysis was conducted using FlowJo^™^ software. Uptake and transfection were evaluated based on the median fluorescence intensity (MFI) of DiR⁺ and eGFP⁺ populations, respectively, from gated histograms.

#### 2.8.2. Live cell imaging

To investigate the uptake and protein expression kinetics of eGFP mRNA LNP, HEK293T and HeLa cells were monitored using the Incucyte^®^ SX5 Live-Cell Analysis System (Sartorius AG, Germany). Briefly, 7,000 HEK293T cells and 5,000 HeLa cells were seeded per well in 96-well plates (Falcon/Corning, Corning, NY) and allowed to adhere for 24 hours. The medium was then removed and replaced with 100 µl of fresh medium containing DiR-labeled eGFP mRNA LNP, resulting in a final dose of 100 ng mRNA per well. The same experimental groups and controls described in Section 2.8.1 were used for consistency and comparative analysis. Both LNP uptake and eGFP mRNA expression were monitored for 48 h. Images were automatically acquired every 4 hours using a 20x objective in both phase-contrast and fluorescence channels. Four distinct images were captured per well throughout the entire time course. Image analysis was performed using the Incucyte^®^ software (Sartorius AG, Germany). The resulting data were exported and subsequently imported into GraphPad Prism for graphing and statistical analysis.

### 2.9. Statistical analysis

Statistical analyses were performed using GraphPad Prism version 10.6.1 (GraphPad Software, USA). Data were analyzed by two-way ANOVA followed by Tukey’s test for multiple comparisons. Statistical significance was defined as *P < 0.05, **P < 0.01, and ***P < 0.001.

## 3. Results

### 3.1. Tuning of manufacturing parameters affects LNP physicochemical properties

To investigate the impact of formulation and process parameters on LNP shape, we implemented a DoE approach by tuning DSPC content (10–70 mol%), lipid-to-RNA ratios (N:P ratio; 3–12), and flow rate ratios (FRR; 3–5). The total flow rate and initial lipid concentration were kept constant at 20 ml/min and 12 mM, respectively.

LNP were prepared via T-junction mixing, in which an ethanol phase containing MC ionizable lipid, DSPC, cholesterol, and DMG-PEG2000 was combined with an aqueous poly A solution at defined ratios. Detailed formulation conditions and composition are listed in **Table S1** and **Table S2**, respectively. By employing DLS and NTA, and generating surface plots, it was identified that increasing DSPC content was the primary factor affecting size, dispersity, and particle concentration, while neither FRR nor N:P ratio had a significant effect within the tested range (**Fig. 1A**).

**Figure 1.**
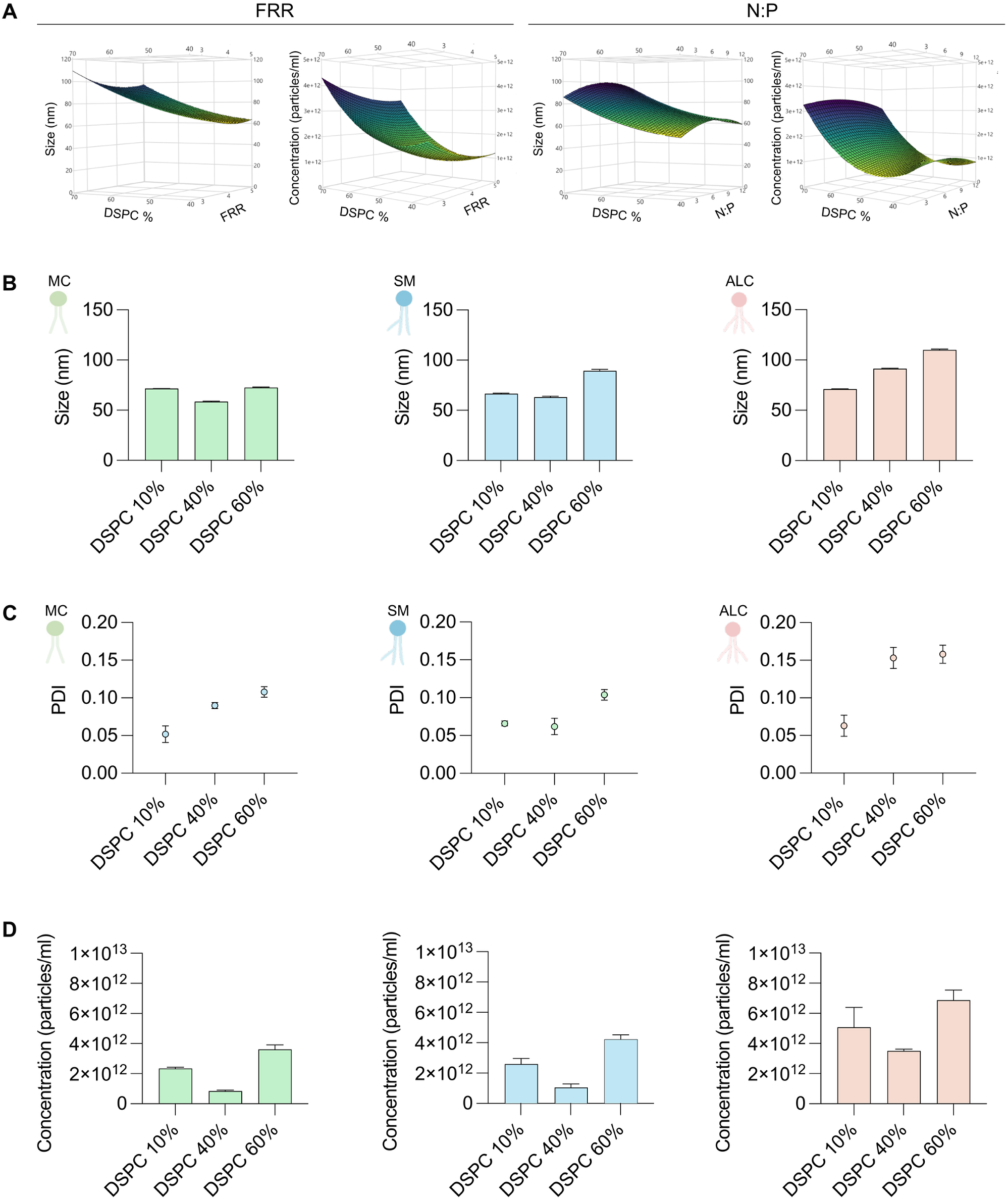
Identification of manufacturing parameters that impact the LNP physicochemical characteristics. (**A**) Response surface plots generated from DoE illustrating the influence of formulation parameters, including DSPC mol%, FRR, and N:P ratio on LNP size, PDI, and particle concentration (particles/ml). Increasing DSPC mol% resulted in larger particles and higher particle concentrations. Since a fixed lipid amount was used, the increase in particle number suggests reduced lipid content per particle, consistent with changes in particle morphology. (**B-D**) Physicochemical characterization of selected LNP formulations spanning the design space and prepared with different ionizable lipids. LNP (**B**) size, (**C**) PDI, and (**D**) concentration. Across all ionizable lipids, increasing DSPC content produced larger particle size and higher particle number, an effect consistently observed across all formulations. Data represent LNP formulated with different ionizable lipids: MC (green), SM (blue), and ALC (red).

To assess the generalizability of this trend, the experiments were repeated using eGFP mRNA and additional ionizable lipids (i.e., SM and ALC) alongside MC, across a range of DSPC contents (10, 40, and 60 mol%) that span the design space. Our findings revealed that across all ionizable lipids, increasing DSPC content from 10% to 60% was associated with a significant increase in LNP size and dispersity (**Fig. 1B,C**). This observation aligns with previous studies reporting that high levels of DSPC or Egg sphingomyelin (ESM) in LNP formulations resulted in larger particles with increased structural heterogeneity^26^.

NTA measurements revealed a decrease in particle number as DSPC content increased from 10% to 40%, followed by a significant increase at 60% (**Fig. 1D**). As a fixed lipid amount was used, this increase in particle number suggests that fewer lipids are incorporated per 60% DSPC-containing particle, possibly indicating a shift from typical spherical, compact vesicles to more vesicular structures. These patterns were largely conserved across formulations prepared with different clinically approved and widely used ionizable lipids, namely MC, SM, and ALC ^32^. Collectively, these results suggest that structural phospholipids play a significant role in driving changes to LNP morphology.

### 3.2. Increasing structural phospholipid content affects LNP shape and morphology

To assess LNP morphological changes under different manufacturing conditions, cryo-TEM was employed. Cryo-TEM imaging revealed that increasing DSPC content induced pronounced structural changes in LNP.

At 10 mol% DSPC, particles, particularly MC and SM, appeared spherical and relatively uniform, exhibiting largely amorphous internal structures (**Fig. 2A** and **Fig. S1**). This morphology is consistent with the known mRNA-LNP structure, in which the mRNA is complexed with the ionizable lipid to form a dense core, surrounded by a monolayer rich in DSPC and PEG-lipids, with cholesterol distributed throughout. Similar structures have been observed in cryo-TEM and small-angle neutron scattering studies^30,33–36^. LNP formulated with ALC showed a distinct multicompartmental morphology. These particles contained an electron-dense, amorphous region next to a more electron-lucent area with internal structure, enclosed by a lipid bilayer (**Fig. 2A**). This heterogeneous internal organization may be attributed to the lower pKa of ALC, which could result in a higher fraction of neutralized ionizable lipid during formation. Reduced electrostatic interactions at this stage may inhibit complete fusion and reorganization, resulting in multicompartmental particles with both electron-dense and lucent regions. Such architectures have been reported previously^24,37^, in which nanogold labeling identified the electron-lucent area as the main site of encapsulated mRNA^37^.

**Figure 2.**
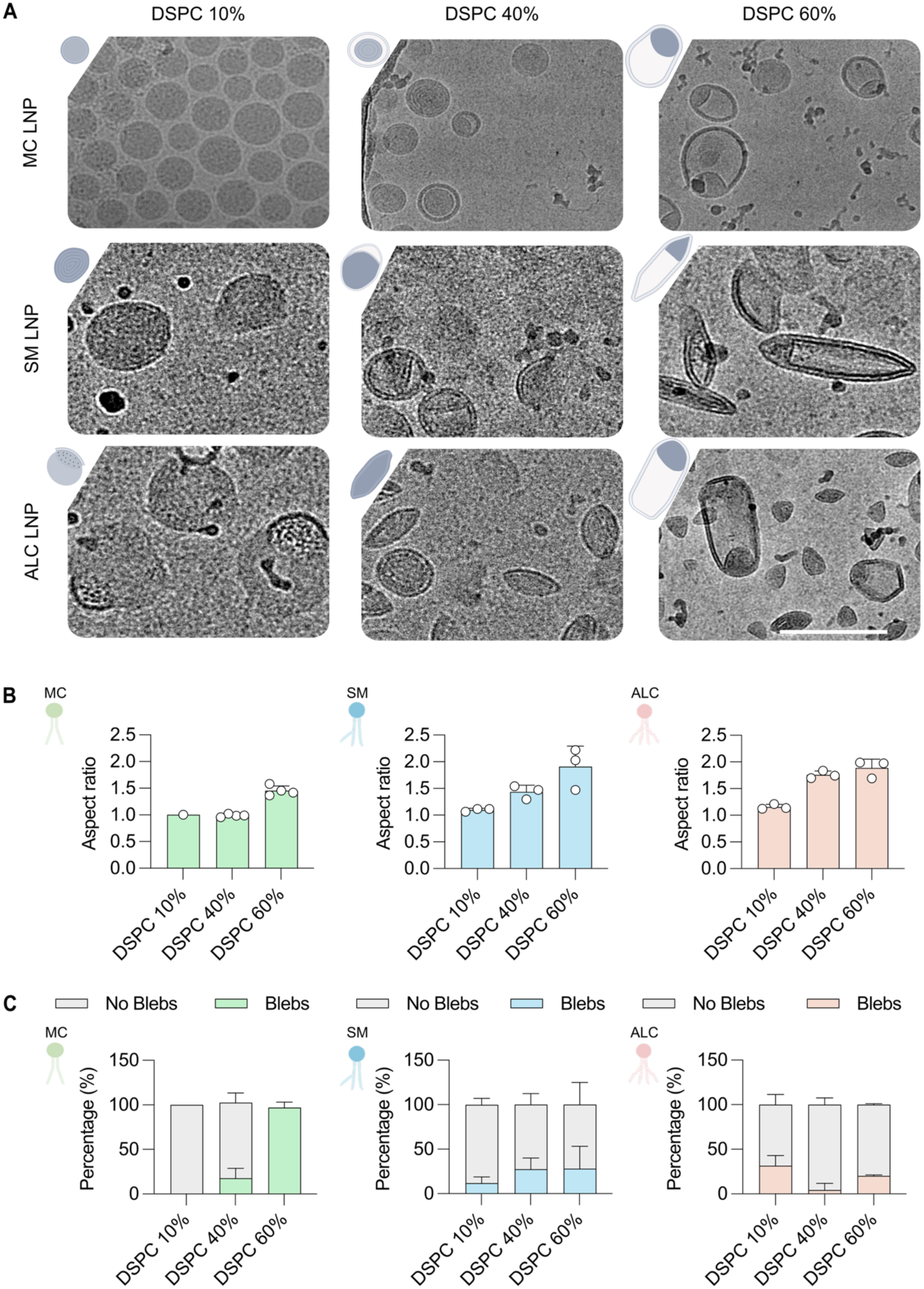
Increasing the percentage of structural lipid (DSPC%) drives a shift in mRNA LNP shape from spherical to non-spherical. (**A**) Cryo-TEM images of eGFP mRNA LNP prepared with different ionizable lipids and increasing percentages of DSPC, demonstrating that changes in formulation composition alter LNP size, shape, and internal structure. Scale 100 nm. (**B**) Aspect ratio of LNP calculated by measuring particle width and height using ImageJ. Each data point represents the average aspect ratio of individual particles measured within a single image; three images were analyzed per formulation. (**C**) Percentage of LNP displaying membrane blebs. Blebbed particles were defined as LNP showing membrane protrusions surrounding an aqueous core in Cryo-TEM images. Quantification was performed by counting blebbed particles within a single image; three representative images were analyzed per formulation.

As seen above, increasing DSPC from 10 to 40 mol% did not result in a significant change in particle size (**Fig. 1B**). However, cryo-TEM revealed the emergence of bilayer features (**Fig. 2A**). This outcome can be explained by examining particle size and concentration: size remained largely unchanged (**Fig. 1B**), whereas particle concentration decreased (**Fig. 1D**), indicating more DSPC was incorporated into existing particles, promoting bilayer formation.

Upon further increasing DSPC content to 60 mol%, LNP exhibited pronounced elongation, adopting balloon-like morphologies for MC and spindle-like morphologies for SM and ALC lipids. These particles displayed an expanded internal aqueous compartment in which the encapsulated mRNA likely resides (**Fig. 2A**).

Quantification of aspect ratio confirmed the qualitative observations, with average aspect ratios increasing from around 1.0 for 10% DSPC to 1.5-2.0 for 60% DSPC for all three ionizable lipids tested (**Fig. 2B**). In addition, upon increasing the DSPC content, bilayer and bleb-like structures became more prevalent, which were particularly pronounced in MC-based LNP, likely due to the chemical structure of the MC ionizable lipid (**Fig. 2C**). These observations indicate that increasing DSPC content induces progressive and reproducible morphological changes, ranging from compact spherical particles to elongated, non-spherical structures.

### 3.3. Increasing structural phospholipid content affects RNA loading

Given the distinct LNP morphologies observed, we evaluated whether these morphological changes affected RNA loading. Bulk analysis of encapsulation efficiency showed that 90 to 95% of RNA, whether for poly A (**Fig. S2A**) or mRNA (**Fig. S2B**), was encapsulated across all formulations. However, single-particle analysis using NanoFCM revealed that the average number of RNA copies per particle declined by approximately 1 mRNA copy when transitioning from 10% to 60% DSPC (**Fig. 3A**). Similarly, increasing DSPC from 10% to 60% led to a decrease in the proportion of RNA-loaded particles by approximately 30 % (**Fig. 3B**). Taking the above information together and considering the particle concentration for 10% and 60% DSPC (**Fig. 1D**), we come to the following realization: low DSPC% results in fewer particles with most particles being loaded; while high DSPC% results in more particles with about half of them being loaded. In addition, low DSPC% particles contain, on average, approximately one additional mRNA copy per particle than high DSPC% particles.

**Figure 3.**
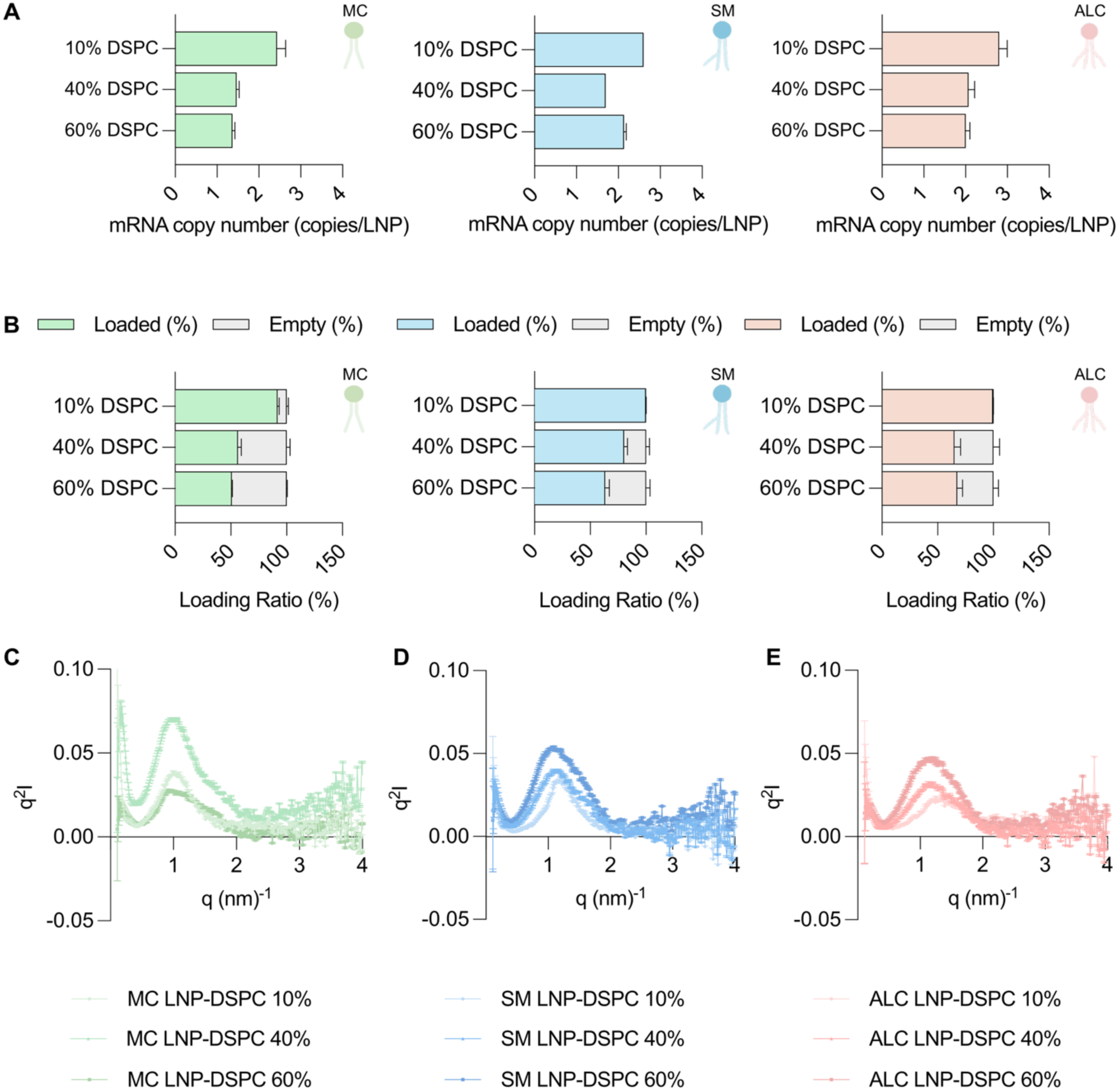
Single particle analysis and SAXS reveal that increasing DSPC content reduces RNA loading and alters LNP structure. (**A-B**) Quantification of RNA loading at the single particle level using NanoFCM following staining of LNP with SYTO 9, a membrane-permeable nucleic acid dye. Flow cytometric analysis was used to determine (**A**) the number of RNA copies per LNP and (**B**) the proportion of RNA-loaded versus empty LNP. Increasing DSPC content within the formulation reduced the number of RNA copies per particle and increased the percentage of empty LNP. (**C-E**) SAXS analysis was performed to assess LNP shape and internal structure. SAXS scattering curves of LNP formulated with (**C**) MC, (**D**) SM, and (**E**) ALC as ionizable lipids, showing DSPC-dependent changes in the low-q and high-q regions, which are associated with particle shape and internal structure, respectively.

SAXS analysis reinforced these findings, revealing DSPC-dependent changes in both particle shape and internal structure (**Fig. 3C-E**). In SAXS, the scattering vector (q) represents the scattering angle. Features observed at low q values (small angles) provide information regarding the overall size and shape of particles, whereas features at high q values (large angles) indicate details about internal structure within the particle.

Considering low q (associated with particle shape), for MC LNP with 10 % DSPC, the SAXS data are well described by a spherical form factor with a radius of ∼ 17 nm. Increasing DSPC content to 40 mol% reduced the fitted radius to 11 nm (**Fig. 3C** and **Fig. S3**), which is consistent with the DLS data indicating a slight decrease in particle size upon increasing DSPC to 40 mol% (**Fig. 1B**). At 60 mol% DSPC, the scattering profiles could no longer be fitted with a spherical model and instead exhibited a power-law decay of ∼ q^-2.5^ at low q, consistent with non-spherical morphologies (**Fig. 3C**). In the higher q regime (associated with internal structure), MC LNP containing 10 % DSPC displayed a single correlation peak at q = 1.03 nm^-1^. Increasing DSPC content to 40 mol% resulted in the appearance of an additional shoulder between q = 1.5 and 2 nm^-1^. At 60 mol% DSPC, the original peak split into two distinct features while the high-q shoulder persisted. This suggests a change in internal structure and possibly a phase transition. However, as increasing DSPC content also promoted the formation of bilayer-rich structures such as blebs, which scatter in a similar q-range, further assignment of these features was not possible based on SAXS alone.

SM LNP behaved similarly to MC LNP. At 10% DSPC, the low-q regime was consistent with spherical particles, whereas increasing the DSPC content to 40% resulted in smaller, more polydisperse particles. At 60% DSPC, an apparent slope of approximately q = 2.5 was again observed in the low-q regime, indicating similar morphologies to those seen for MC LNP at 60% DSPC (**Fig. 3D**). The presence of bleb-like and/or elongated structures was also observed by Cryo-TEM imaging (**Fig. 2A**). In contrast, ALC-based LNP exhibited distinct behavior. At 10 mol% DSPC, the SAXS profiles lacked a clear Guinier regime, consistent with heterogeneous particle size and/or shape (**Fig. 3E**).

This observation was supported by Cryo-TEM images, which revealed the presence of electron dense and electron-lucent regions within individual particles (**Fig. 2A**). The internal structure of ALC-based LNP showed a single dominant correlation peak that shifted from q = 1.38 nm^-1^ at 10 mol% DSPC to q = 1.2 nm^-1^ at 40 mol% DSPC, corresponding to an increase in correlation length from 4.5 to 5.2 nm. Increasing DSPC content to 40 mol% also resulted in a transition toward smaller, prolate ellipsoidal particles, a morphology that was maintained at 60 mol% DSPC (**Fig. 3E**). Increasing DSPC content further to 60 mol% increased the peak area without altering its position or width, suggesting an increase in the fraction of ordered internal material without a change in domain size; no additional peaks or shoulders were observed (**Fig. 3E**).

### 3.4. Non-spherical LNP enhance eGFP mRNA expression

To examine the influence of LNP morphology and RNA loading on cellular uptake and transfection, HEK293T and HeLa cells, two widely used models for mRNA transfection, were treated with eGFP mRNA LNP. Prior to cell experiments, LNP stability was evaluated in PBS and BSA-supplemented PBS at 37 °C for 24 h. DLS measurements confirmed stable particle size in both media over the incubation period (**Fig. S4** and **Fig. S5**). Following stability assessment, eGFP mRNA LNP were added to HEK293T and HeLa cells, and after 24 h, LNP uptake (DiR^+^ cells) as well as eGFP expression (eGFP^+^ cells) were quantified via flow cytometry by gating on live single cells (**Fig. 4A,B**). To enable direct comparison, cells were treated with LNP formulations delivering an equal amount of eGFP mRNA (200 ng).

**Figure 4.**
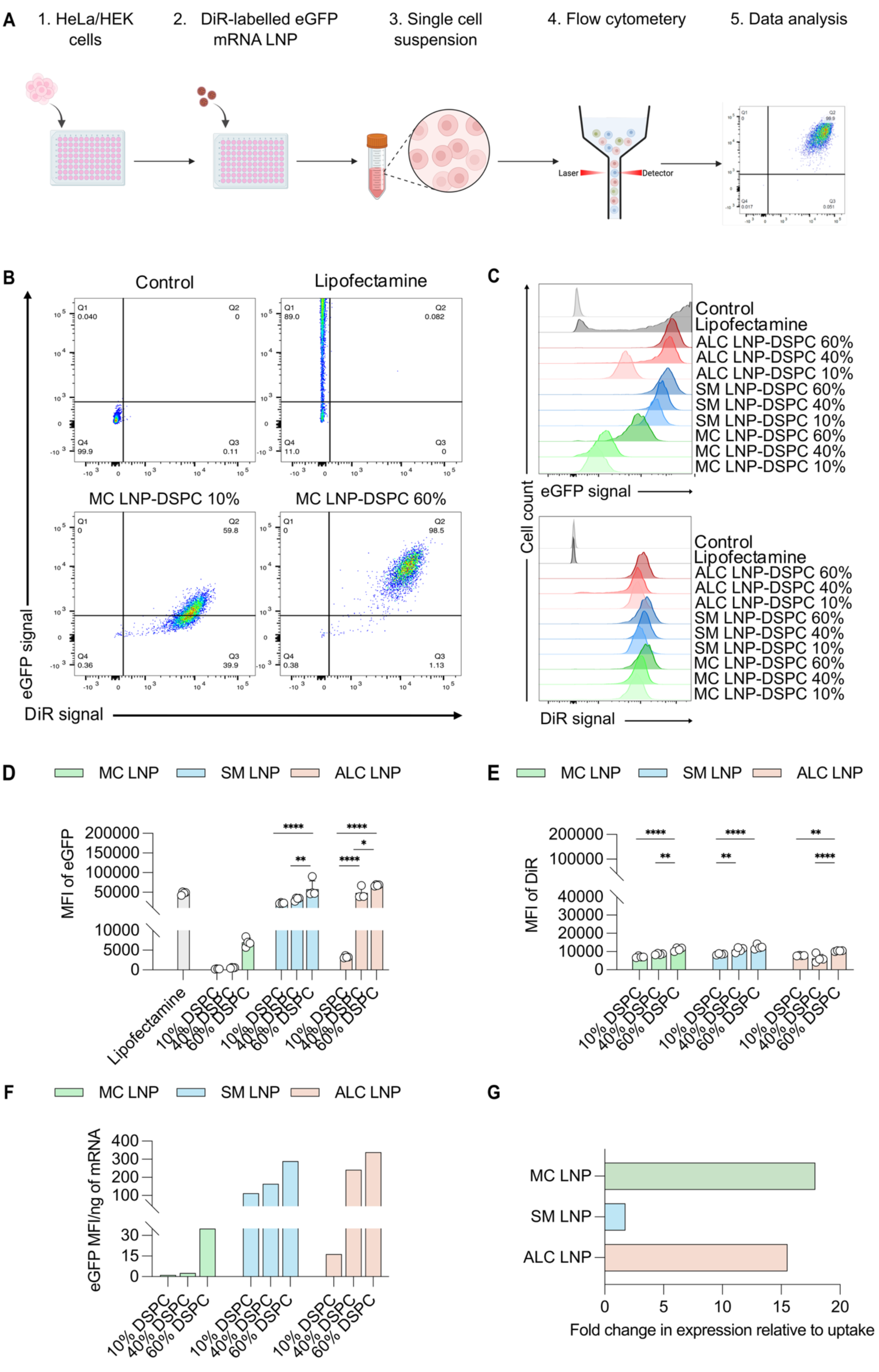
Increasing DSPC content shifts LNP shape and enhances cellular uptake and eGFP mRNA expression across all tested ionizable lipids. (**A**) Schematic illustrating key experimental steps for assessing LNP uptake and eGFP mRNA expression. (**B**) Gating strategy used for analysis of LNP-treated cells. (**C**) Representative histograms showing expression of eGFP mRNA (eGFP^+^) and uptake (DiR^+^) for LNP formulated with varying DSPC percentages. (**D–E**) Quantification of mean fluorescence intensity (MFI) showing that increasing DSPC enhances eGFP expression and LNP uptake. (**D**) MFI of eGFP in HEK cells following eGFP mRNA delivery by LNP (**E**) MFI of DiR signal representing LNP uptake in HEK cells. (**F)** eGFP expression normalized to mRNA levels to evaluate expression efficiency, showing higher efficiency as DSPC content increased. (**G**) Fold increase in eGFP expression normalized to DiR uptake comparing standard spherical LNP with 10% DSPC to non-spherical LNP with 60% DSPC, showing higher normalized expression for non-spherical, high-DSPC LNP. Results are presented as mean ± SD (n=4). P values are indicated as * < 0.05, ** < 0.01, *** < 0.001, and **** < 0.0001.

The analysis revealed an inverse relationship between sphericity and expression, as the non-spherical LNP formed with 60% DSPC resulted in higher eGFP expression than spherical LNP across both cell lines (**Fig. 4C,D** and **Fig. S6A**). The magnitude of this enhancement depended on the ionizable lipid used. Overall, it appeared that MC-based formulations exhibited only a modest increase in expression when compared to SM-and ALC-based LPN that reached substantially higher absolute expression levels (**Fig. 4D**). However, within each ionizable lipid group, the response to increasing DSPC differed. MC-based LNP showed the greatest relative improvement, with minimal expression at 10% and 40% DSPC, and a marked increase at 60% DSPC. In contrast, SM- and ALC-based LNP already exhibited higher expression at lower DSPC content, with a more moderate increase in expression as DSPC content was raised (**Fig. 4D** and **Fig. S6A**).

To assess whether the observed enhancement in transfection was due to increased nanoparticle uptake, flow cytometry was used to quantify uptake of fluorescent LNP. Uptake was increased for non-spherical formulations (60% DSPC) compared to spherical (10% DSPC) by approximately 1.5-fold (**Fig. 4E** and **Fig. S6B**). The higher uptake observed for non-spherical LNP is consistent with their larger size and elongated geometry, which increase surface area and promote more efficient interactions with the cell membrane ^28,38–40^. However, the higher uptake should also be viewed in the light of the difference in formulation between the 10 and 60% LNP. The non-spherical LNP contained fewer mRNA copies per particle, and a higher proportion of empty particles. Additionally, since an equal amount of DiR was used to label both formulations, the larger number of particles also resulted in less DiR per individual 60 mol% DSPC LNP (**Fig. 1D**, **Fig. 3A,B**, **Fig. 4D,E**, and **Fig. S7**). Therefore, approximately four times more particles of the 60% LNP were required to deliver the same mRNA dose as the 10 mol% DSPC LNP. Notably, the empty particles in 60% LNP formulations contribute to the measured DiR uptake but do not contribute to mRNA delivery. Despite this, the non-spherical LNP achieved substantially higher eGFP expression.

Normalization of expression to mRNA amount (**Fig. 4F**) and to uptake (**Fig. 4G**) still revealed a substantial improvement in expression for the non-spherical LNP. These findings suggest that increased cellular uptake contributes to the enhanced transfection of non-spherical LNP but is not sufficient to explain the magnitude of the observed improvement. Rather, the sustained improvement after normalization to uptake points toward an additional contribution from LNP morphology and internal architecture to functional mRNA delivery.

### 3.5. Non-spherical LNP exhibit sustained eGFP expression over time

Live-cell imaging was performed over 48 h to investigate the differences in eGFP mRNA expression kinetics between LNP with different morphologies (**Fig. 5A,B** and **Fig. S8-10**). Time-resolved analysis revealed distinct expression profiles between spherical and non-spherical LNP. Spherical LNP (10% DSPC) displayed a classical pattern, with an initial increase in expression, followed by a gradual reduction in the rate of increase, and a plateau at later time points. In contrast, non-spherical LNP (60% DSPC) exhibited continuous increases in expression throughout the 48 h period (**Fig. 5C** and **Fig. S8-10**). At 24 h, non-spherical LNP showed an approximately 3.5-fold increase in eGFP expression compared to spherical LNP, which increased to 12-fold by 48 h (**Fig. 5C**). Of note, all LNP formulations were dosed to deliver equivalent total mRNA amount; therefore, the higher expression observed with 60% DSPC likely reflects an increased expression efficiency.

**Figure 5.**
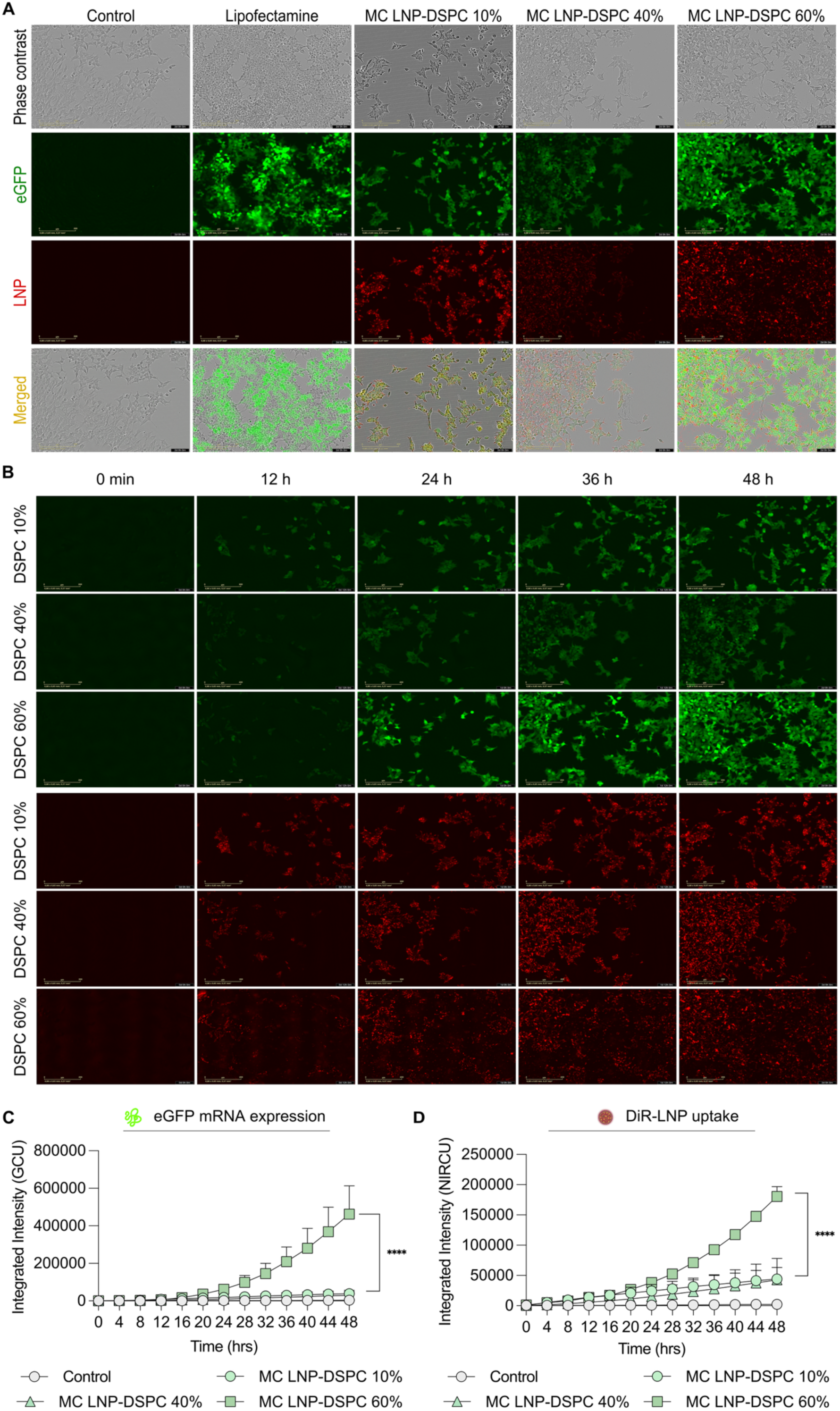
Non-spherical LNP containing a high percentage of structural lipid (DSPC) exhibit enhanced cellular uptake and prolonged eGFP expression in HEK cells. (**A**) Fluorescence microscopy images showing LNP uptake (DiR) and eGFP expression in HEK cells treated with MC-based LNP containing increasing DSPC content after 48 h of incubation. Scale bar: 200 µm. (**B**) Time-lapse fluorescence microscopy images acquired at 0, 12, 24, 32, and 48 h, illustrating the changes in eGFP expression and LNP uptake over time. (**C-D**) Quantification of integrated intensity over 48 h, measured at 4 h intervals. Four distinct images were captured per well at each time point using the Incucyte^®^ software (Sartorius AG, Germany). (**C**) Quantification of eGFP expression, measured as green channel unit (GCU), showing changes in expression over 48 h. (**D**) Quantification of DiR-labelled LNP uptake, measured as near-infrared channel unit (NIRCU), showing changes in LNP uptake over time. Results are presented as mean ± SD (n=3). P values are indicated as * < 0.05, ** < 0.01, *** < 0.001, and **** < 0.0001.

A similar trend was observed for LNP uptake. Non-spherical LNP showed a 1.5-fold increase in uptake at 24 h and a 4-fold increase at 48 h compared to spherical LNP, indicating that the non-spherical LNP prepared with high DSPC mol% were taken up more efficiently than spherical LNP (**Fig. 5D** and **Fig. S7**). Similar to the flow cytometry experiment, a larger number of particles had to be delivered to obtain the same amount of RNA, which explains the increased nanoparticle uptake signal observed for non-spherical particles (**Fig. S7**). These patterns were consistent for both cell lines across LNP formulated with different ionizable lipids (**Fig. S8-10**).

The agreement in results between flow cytometry and live cell imaging indicates that the enhanced transfection efficiency of non-spherical LNP is at least partly related to improved cellular interaction. However, the substantially greater increase in protein expression relative to uptake (**Fig. S7**) suggests that particle shape plays a significant role in this process.

## 4. Discussion

This work set out to understand which composition and manufacturing parameters influence LNP morphology, and how these structural differences affect LNP potency. Recent studies have suggested that the morphology and internal architecture of LNP particles can influence biological activity by affecting cellular uptake, transfection efficiency, and even biodistribution^17,22,26–28,41,42^. Here, we provide a deeper, mechanistic assessment demonstrating that controlled modulation of lipid composition, particularly the fraction of the helper structural lipid DSPC, drives defined architectural changes in LNP that translate into distinct and predictable differences in biological performance.

To systematically dissect these effects, we generated an LNP library by varying lipid composition, FRR, and N:P ratio. Among these factors, lipid composition, specifically the molar fraction of DSPC, proved to be the main determinant of particle characteristics, influencing size, polydispersity, and particle concentration. All formulations exhibited the same overall trend: increasing DSPC content produced larger particles, and higher particle counts per mole of lipid (indicating fewer lipid molecules per particle), suggesting a more vesicular shape. Notably, these changes were accompanied by a reduction in RNA-loaded particles (i.e., a larger fraction of empty particles) and lower RNA copy per particle. Similar trends have been reported, where increasing helper-lipid content (ESM or DSPC) results in larger and higher numbers of LNP compared to the standard Onpattro^™^ formulation with lower helper lipid fractions^26^. However, these studies assumed uniform RNA encapsulation across particles, attributing reduced RNA per particle solely to redistribution across a larger particle population. Our single-particle analysis refines this interpretation, revealing that higher helper-lipid content not only produces more particles but also generates a substantial fraction of empty ones.

The concurrent increase in particle size and number with increasing DSPC content already suggests significant changes in morphology, as it implies that fewer lipid molecules constitute a single particle. This means that the particles not only expand but also contain a substantial aqueous volume. Cryo-TEM imaging confirmed that the observed physicochemical differences translated into distinct morphological changes across formulations. These structural transitions can be understood from both structural and thermodynamic perspectives. Since DSPC is a high transition temperature, fully saturated phospholipid, it naturally tends to form rigid lamellar bilayers. At lower concentrations (∼10%), it mainly stabilizes the particle surface, leading to compact, spherical LNP^30,43,44^, whereas higher fractions promote internal bilayer domains and increased water incorporation. Beyond a critical threshold (∼30-40%), which most likely depends on the phospholipid used^26^, continued bilayer growth leads to elongated morphologies, consistent with reports for ESM-containing LNP^26,45^.

These structural changes are further influenced by the chemical structure of the ionizable lipid. ALC-based LNP exhibited larger sizes and earlier onset of bleb-like morphologies, consistent with the lipid’s larger cross-sectional area and tendency toward phase segregation^46,47^. In contrast, SM-based LNP, with its asymmetrical structure and phospholipid-like packing, favored more ordered bilayers, whereas MC displayed intermediate behavior^46^. The influence of DSPC percentage and ionizable lipid chemistry on LNP shape was further corroborated by SAXS analysis. In the low-q regime, the scattering profiles revealed a transition from compact spherical particles to blebbed or elongated morphologies as DSPC content increased.

These findings naturally raise the question of how such structural alterations, and the accompanying reduction in RNA loading, affect the biological activity of eGFP mRNA LNP, particularly their delivery and expression efficiency. To address this, HEK293T and HeLa cells were treated with LNP of different morphologies, and both uptake and protein expression were quantified after 24 hours using flow cytometry. Non-spherical LNP consistently showed higher eGFP expression compared to spherical LNP. Live-cell imaging further revealed that non-spherical LNP exhibited a distinct, sustained expression profile compared to spherical ones. Notably, the magnitude of this shape-dependent enhancement in expression varied with the ionizable lipid. Several factors could explain this improved performance, including variations in cellular uptake, LNP shape, internal structure, and RNA loading.

First, increased cellular uptake may partly contribute to the enhanced transfection efficiency of non-spherical LNP, presumably due to their unique surface geometry, which creates multiple membrane contact points that favor wrapping and internalization, or to the higher rate of endocytosis commonly observed in cancer cells^38–40,48,49^. However, cellular entry alone does not necessarily translate into functional mRNA delivery, as LNP transfection is typically constrained by downstream intracellular barriers, most notably endosomal escape^43,50,51^. Therefore, formulation features that facilitate mRNA release into the cytosol may become increasingly important. In this context, the internal structure of LNP becomes important, with topological features that increase lipid–water interfacial area having been proposed to facilitate membrane interactions and RNA delivery into the cytosol^45,51^. The bilayer-rich, blebbed structures seen in non-spherical LNP might therefore offer structural features that promote these interactions and aid in cytosolic RNA delivery. However, the contribution of these structural features may also depend on the ionizable lipid. Previous studies have shown that the chemical structure and pKa of the ionizable lipid influence both endosomal escape efficiency and membrane fusion behavior, thereby affecting the amount of mRNA that reaches the cytosol for translation^52,53^. Differences in these characteristics could explain the variation in expression observed among ionizable lipids^44,50^.

Beyond internal structure and ionizable lipid chemistry, RNA loading may further contribute to the observed enhancement in transfection. Interestingly, recent studies have shown that LNP carrying higher mRNA amounts often have lower transfection efficiency^28,54^, as excessive mRNA loading can shift particles away from an optimal N:P ratio, altering internal structure and potentially hindering endosomal escape. It appears that particles with low to moderate loading may deliver better results by maintaining internal structures that are more conducive to membrane interaction and release. In this regard, the improved performance of non-spherical LNP despite carrying less mRNA aligns with a growing understanding that optimal, rather than maximum, payload loading drives effective delivery and expression. In addition to LNP internal structure and RNA loading, particle morphology may influence LNP intracellular trafficking. Non-spherical LNP have been associated with prolonged intracellular retention, reduced exocytic recycling, and delayed degradation, providing more opportunities for endosomal escape ^22,39,40,55^. Finally, the vesicular or partially aqueous internal core of non-spherical LNP may protect mRNA from lipid degradation ^26,45,56^, thus contributing to sustained protein expression^26,45^.

In conclusion, by precisely controlling lipid composition, it is possible to generate LNP with varied external morphologies and internal structures. These structural differences influence how mRNA is distributed in LNP at the single-particle level, ultimately affecting transfection efficiency and protein expression kinetics. Overall, our findings expand the understanding of the LNP shape–function relationship and highlight morphology, together with internal structure and RNA loading, as functionally relevant design variables that should be integrated into the rational design of LNP for improved mRNA delivery and expression.

## 5. Conclusions

In this study, we aimed to elucidate how process and formulation parameters influence the morphology and performance of mRNA LNP. By systematically varying the structural lipid content and the type of ionizable lipid, we uncovered clear relationships among lipid composition, particle morphology, and biological function.

Cryo-TEM and SAXS analyses revealed that increasing DSPC content drives a transition from compact spherical to elongated or non-spherical LNP. These structural shifts were found to influence RNA loading and significantly improve cellular uptake and expression kinetics.

While spherical particles remain the benchmark for clinical formulations, our study establishes that non-spherical, elongated LNP can mediate superior, more durable expression despite reduced RNA loading. These findings refine our understanding of LNP shape–function relationships and highlight morphology as a tunable design parameter for improving mRNA therapeutics.

Future work should investigate how LNP structure influences biodistribution and transfection in vivo, as such insights will enable the rational design of delivery systems that combine well-defined particle shape with efficient, sustained mRNA delivery.

## Data availability

The data are available from the authors upon request.

## Author contribution

AMS and JM conceptualized and supervised the study. SE and AMS designed the experiments. SE executed all experiments involving LNP preparation, characterization, cellular uptake, and transfection studies. SE and SS prepared and imaged Cryo-TEM samples. BP and JB conducted single-particle LNP analyses using NanoFCM. SE and MW performed NTA measurements. TH designed and provided the GFP mRNA plasmid construct. SE carried out in-house IVT of mRNA. SE performed statistical analyses. SE and AMS designed the figures. SE drafted the manuscript. AMS, JM, and TL acquired funding. All authors contributed to data interpretation, manuscript revision, and approved the final version for submission.

## Acknowledgements

The authors acknowledge financial support from the German Research Foundation (DFG; GRK 2375 Tumor-Targeted Drug Delivery (331065168) to SE, TH, JMM, TL, and Excellence Initiative RWTH JPI 2021 to AMS). The flow cytometry measurements were supported by the Flow Cytometry Facility, a core facility of the Interdisciplinary Center for Clinical Research (IZKF) Aachen within the Faculty of Medicine at RWTH Aachen University. Authors acknowledge the electron microscopy training, imaging, and access time granted by the life science Electron microscopic facility of the Ernst-Ruska Centre (ERC-3) at Forschungszentrum Jülich (FZJ), Germany. Schematics were created with BioRender.com.

## Notes

Ben Peacock and Joseph Brealey are employed by NanoFCM Co., Ltd., Nottingham, UK.

