## Supplementary Material for "Lipid nanoparticle shape and morphology determine the magnitude and duration of mRNA transfection"

**Table S1.** Summary of experimental runs generated by DoE software and the corresponding output parameters, including LNP size, PDI, and particle concentration.

| Run | Lipid:RNA | FRR | DSPC % | Size (nm) | PDI | Concentration (particles/ml) |
| --- | --- | --- | --- | --- | --- | --- |
| 1 | 6 | 4 | 60 | 82,52 | 0,148 | 1,78E+12 |
| 2 | 3 | 3 | 40 | 67,46 | 0,114 | 1,33E+12 |
| 3 | 6 | 5 | 40 | 66,23 | 0,166 | 7,66E+11 |
| 4 | 3 | 4 | 70 | 93,69 | 0,095 | 3,00E+12 |
| 5 | 12 | 3 | 60 | 83,66 | 0,12 | 2,87E+12 |
| 6 | 12 | 4 | 40 | 59,97 | 0,064 | 1,28E+12 |
| 7 | 6 | 3 | 70 | 107,8 | 0,1 | 4,39E+12 |
| 8 | 3 | 5 | 50 | 69,52 | 0,111 | 1,63E+12 |
| 9 | 12 | 5 | 70 | 96,41 | 0,087 | 2,79E+12 |
| 10 | 12 | 3 | 50 | 72,72 | 0,082 | 1460000000 |

**Table S2.** Summary of the lipid components and their molar ratios for the different LNP formulations used in this study.

| Lipid composition | Molar Percentage (%) |  |  |
| --- | --- | --- | --- |
|  | 10% DSPC | 40% DSPC | 60% DSPC |
| Ionizable lipid | 50 | 33 | 21.5 |
| Phospholipid | 10 | 40 | 60 |
| Cholesterol | 38.5 | 25.5 | 17 |
| PEG lipid | 1.5 | 1.5 | 1.5 |

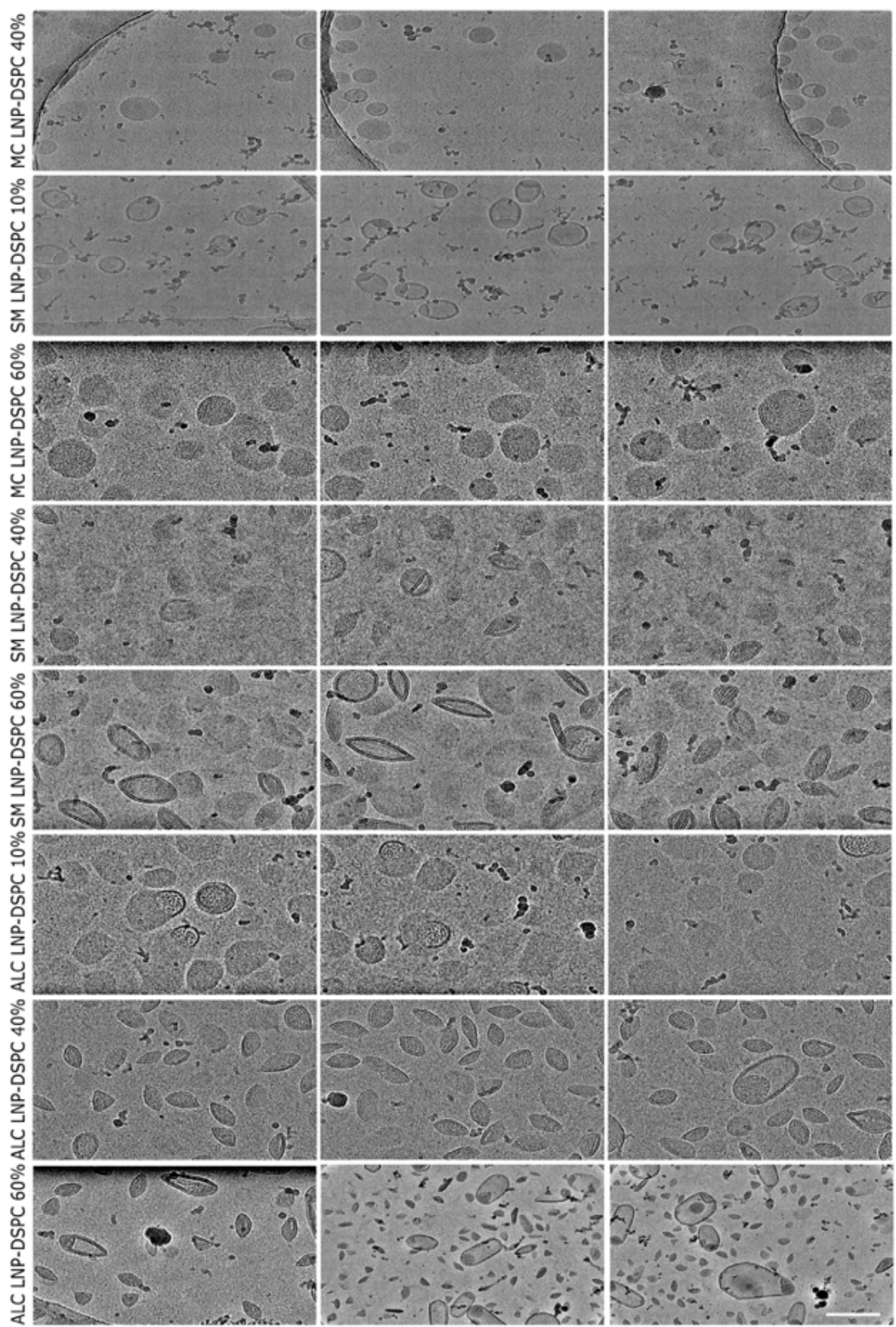

**Figure S1. Representative CryoTEM images of the LNP.** Three representative micrographs are shown for each formulation to illustrate particle morphology and sample consistency. Images were acquired from a single LNP preparation. Scale bar = 100 nm.

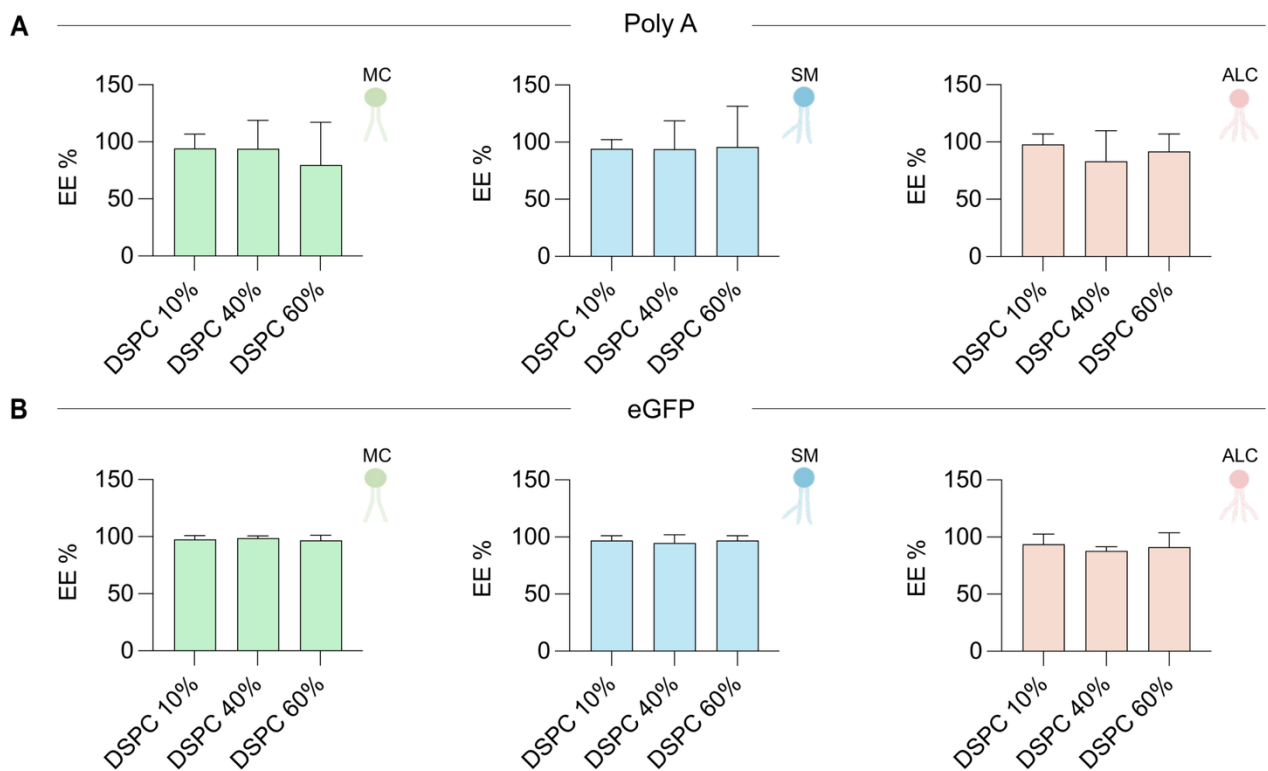

**Figure S2. High RNA encapsulation efficiency is maintained across DSPC contents, ionizable lipids, and RNA payloads** (A) Encapsulation efficiency of mock RNA, Poly A, across formulations with varying DSPC content and ionizable lipid type. (B) Encapsulation efficiency of eGFP mRNA under the same formulation conditions. Data are presented as mean ± SD (n = 3).

### SAXS model for fitting the sphere-like LNP

$$I(q) = A_{sph} \int_0^\infty p(R) (\Delta\rho)^2 V(R)^2 P(q, R) dR + A_{peak} e^{-\frac{(q-q_0)^2}{2\sigma_q^2}} + B$$

$$P(q, R) = \left[ \frac{3(\sin(qR) - qR \cos(qR))}{(qR)^3} \right]^2$$

$$p(R) = \frac{1}{\sqrt{(2\pi)\sigma_R}} e^{-\frac{(R-R_0)^2}{2\sigma_R^2}}$$

$$V(R) = \frac{4\pi R^3}{3}$$

$$\sigma_R = PD R_0$$

The scattering intensity was modeled as the sum of a polydisperse spherical form factor and an empirical Gaussian peak accounting for correlations within the LNP interior. The particle radii were assumed to follow a Gaussian distribution with mean radius $R_0$ and standard deviation $\sigma_R = PD R_0$. The Gaussian peak was characterized by its amplitude $A_{peak}$, peak position $q_0$, and width $\sigma_q$. A constant incoherent background $B$ was additionally included. The spherical contribution was normalized such that $I_0$ represents its zero-angle intensity. The peak here was fitted to account for its contribution in the $q$ region for the spherical form factor. The contrast was set to 1, and the $A_{sphere}$ parameter was left free. This model was implemented in Python and uses the package jscatter.

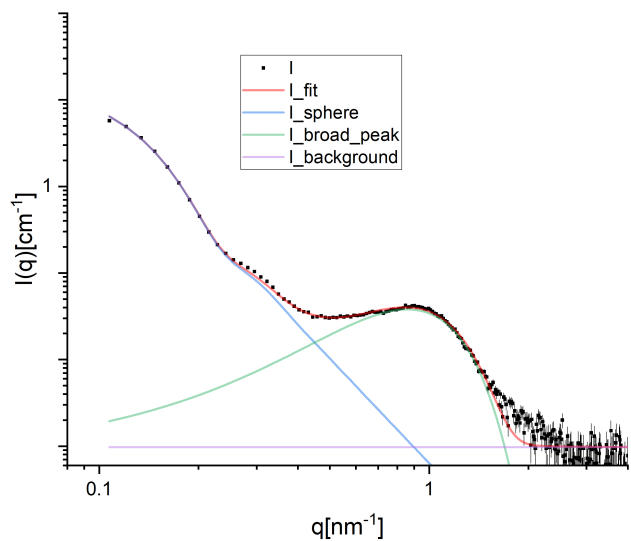

**Figure S3. Scattering curve of MC LNP-DSPC 10%.** Representative fit for the curves showing low $q$ oscillations and mostly spherical LNP in the cryo-TEM. The full fit as well as the fit components defining the model described above. The peak positions described in the main text were fitted with a Gaussian peak function only to the Kratky representations to provide a qualitative description of how peaks primarily affected by internal structures change.

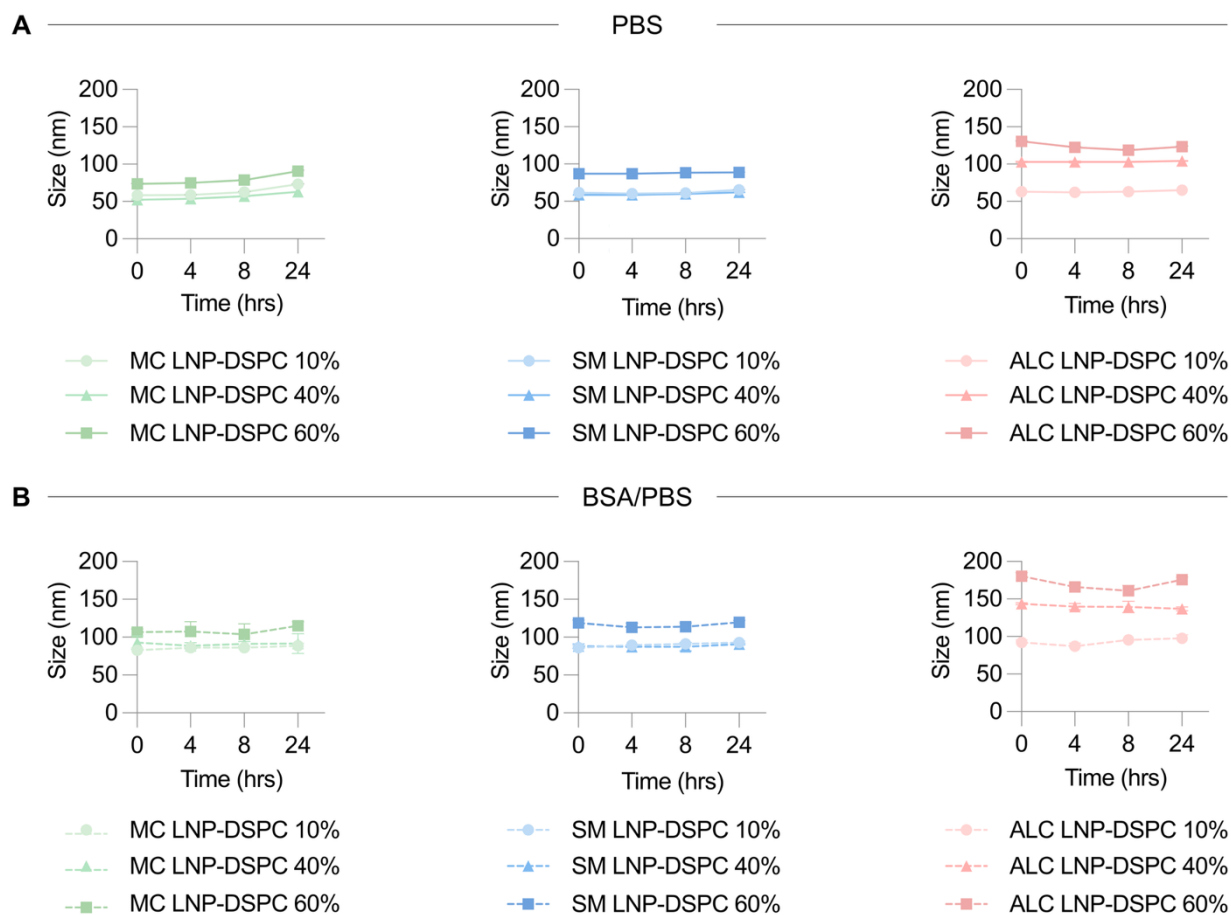

**Figure S4. LNP remain stable over 24 h in PBS and BSA/PBS.** Size of LNP with varying DSPC content, as measured by DLS after 0, 4, 8, and 24 h of incubation at 37 °C under constant shaking in **(A)** PBS and **(B)** BSA/PBS.

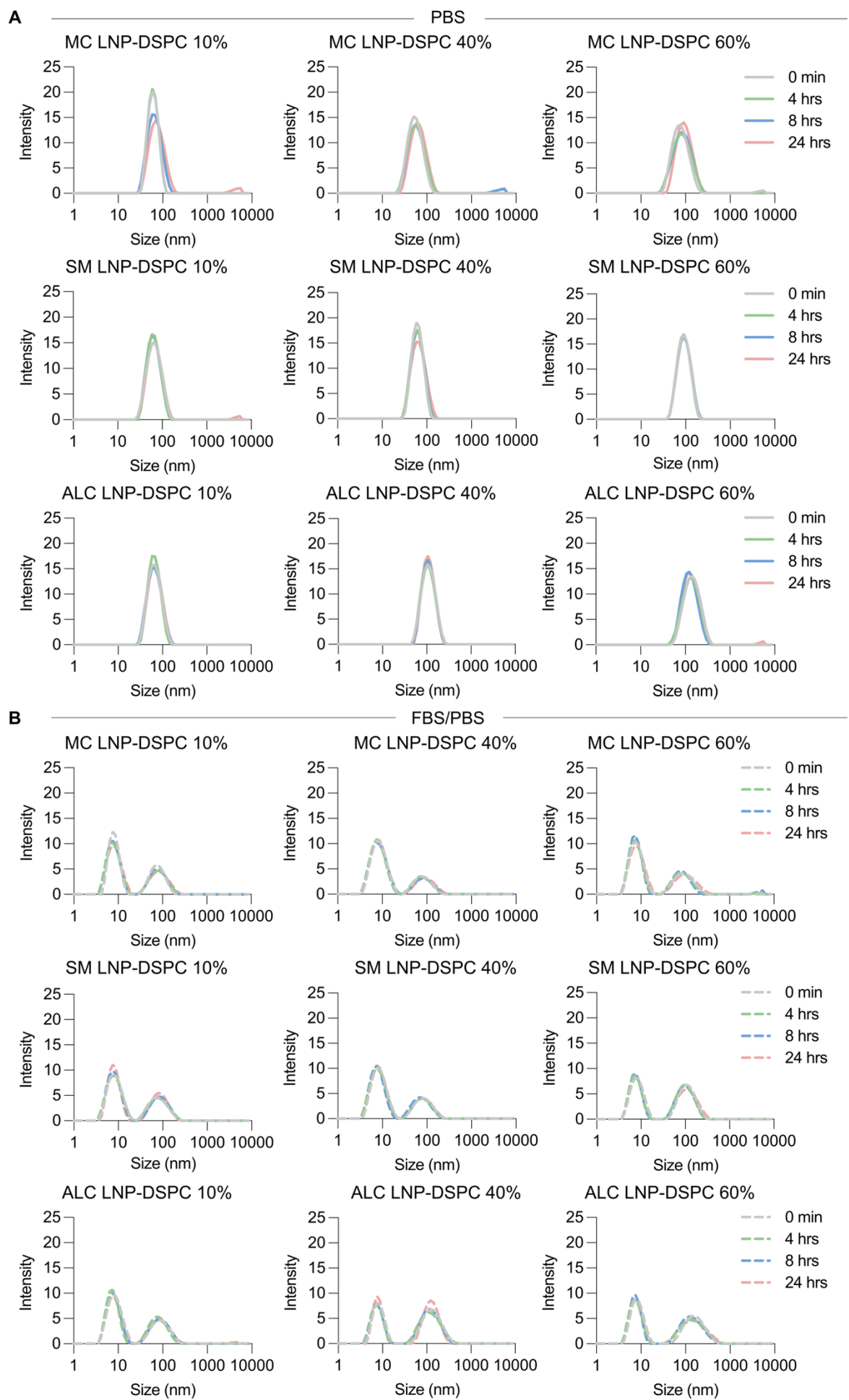

**Figure S5. LNP maintain their size and dispersity over 24 h in PBS and BSA/PBS.** DLS intensity charts of the different LNP formulations after incubation in **(A)** PBS or **(B)** BSA/PBS for 0, 4, 8, and 24 h at 37 °C under constant shaking, showing that LNP maintained stable size distributions over the 24 h period in both media.

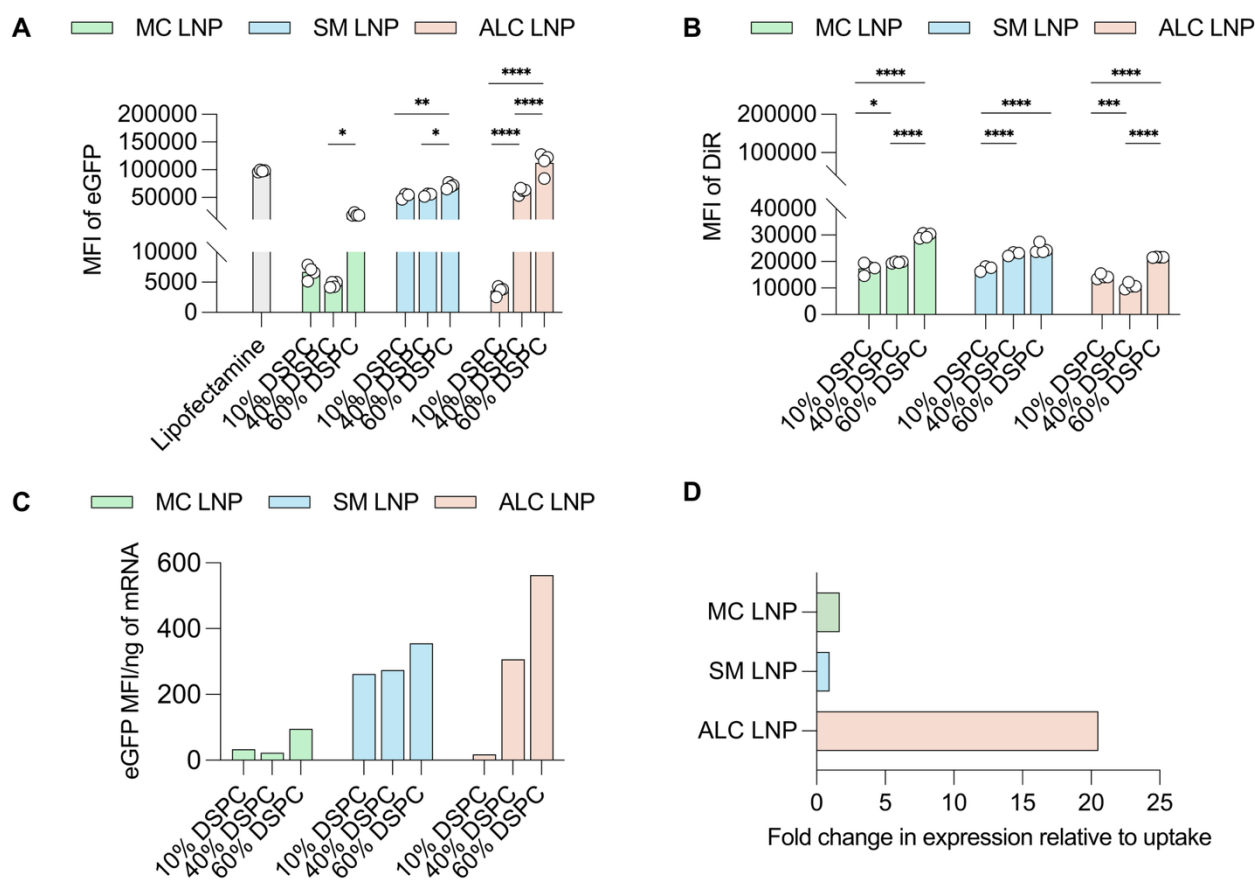

**Figure S6. Quantification of eGFP LNP mRNA uptake and expression formulations.** (A) Mean fluorescence intensity (MFI) of eGFP expression in HeLa cells following mRNA delivery by LNP formulated with different ionizable lipids and DSPC percentages, showing increased eGFP expression with increasing DSPC content. (B) MFI of DiR fluorescence in HeLa cells as a measure of LNP uptake, showing increased cellular uptake with increasing DSPC content. (C) eGFP MFI normalized to RNA amount to assess expression efficiency, showing greater expression per amount of encapsulated RNA for non-spherical, high-DSPC LNP. (D) Fold increase in transfection comparing standard spherical LNP (DSPC 10%) and non-spherical LNP (DSPC 60%).

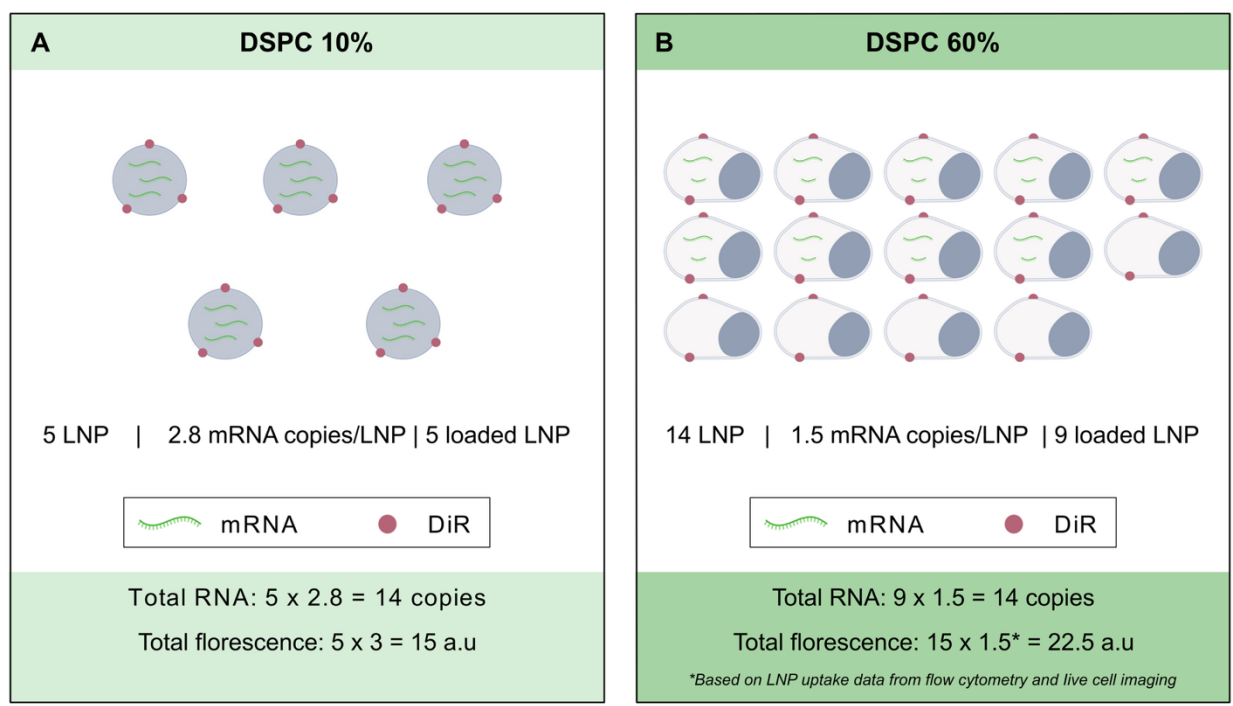

**Figure S7. Schematic depicting the distribution of mRNA and DiR dye among LNP with different morphologies for the delivery of the same mRNA dose, using 14 mRNA copies as an example. (A)** Spherical LNP containing MC ionizable lipid and 10% DSPC have approximately 2.8 RNA copies per particle, as measured by NanoFCM. Therefore, fewer particles are needed to deliver a given RNA amount. In this schematic example, each spherical LNP contains 3 arbitrary units (a.u.) of DiR, representing the relative amount of dye per particle. To deliver approximately 14 RNA copies, around 5 particles are required. This corresponds to a total DiR signal of 15 a.u., calculated as 5 particles × 3 DiR units per particle. **(B)** Non-spherical LNP containing a higher DSPC percentage have a more vesicular structure and have fewer RNA copies per particle (1.5 copies). Therefore, approximately three times more particles are needed to deliver the same total RNA amount. Due to their more vesicular structure, these particles contain more lipid per LNP, and, given that the molar percentage of DiR was kept constant across formulations, these LNP are also expected to contain less DiR per particle. Experimentally, flow cytometry and live-cell imaging showed approximately 1.5-fold higher DiR uptake for the non-spherical LNP. Using the spherical LNP signal of 15 a.u. as a reference, this corresponds to a total DiR signal of approximately 22.5 a.u. ($1.5 \times 15$ a.u.) for non-spherical NP. While increased uptake might partly account for the greater expression of non-spherical LNP, a closer look at the schematic shows that these particles carry less RNA per particle and have a higher proportion of empty particles, which contribute to the measured DiR uptake without delivering mRNA. Nevertheless, they achieve significantly higher eGFP expression, indicating that uptake isn't the sole factor behind the improved expression. Instead, the unique shape and internal structure of non-spherical LNP may promote more efficient functional mRNA delivery following cellular uptake.

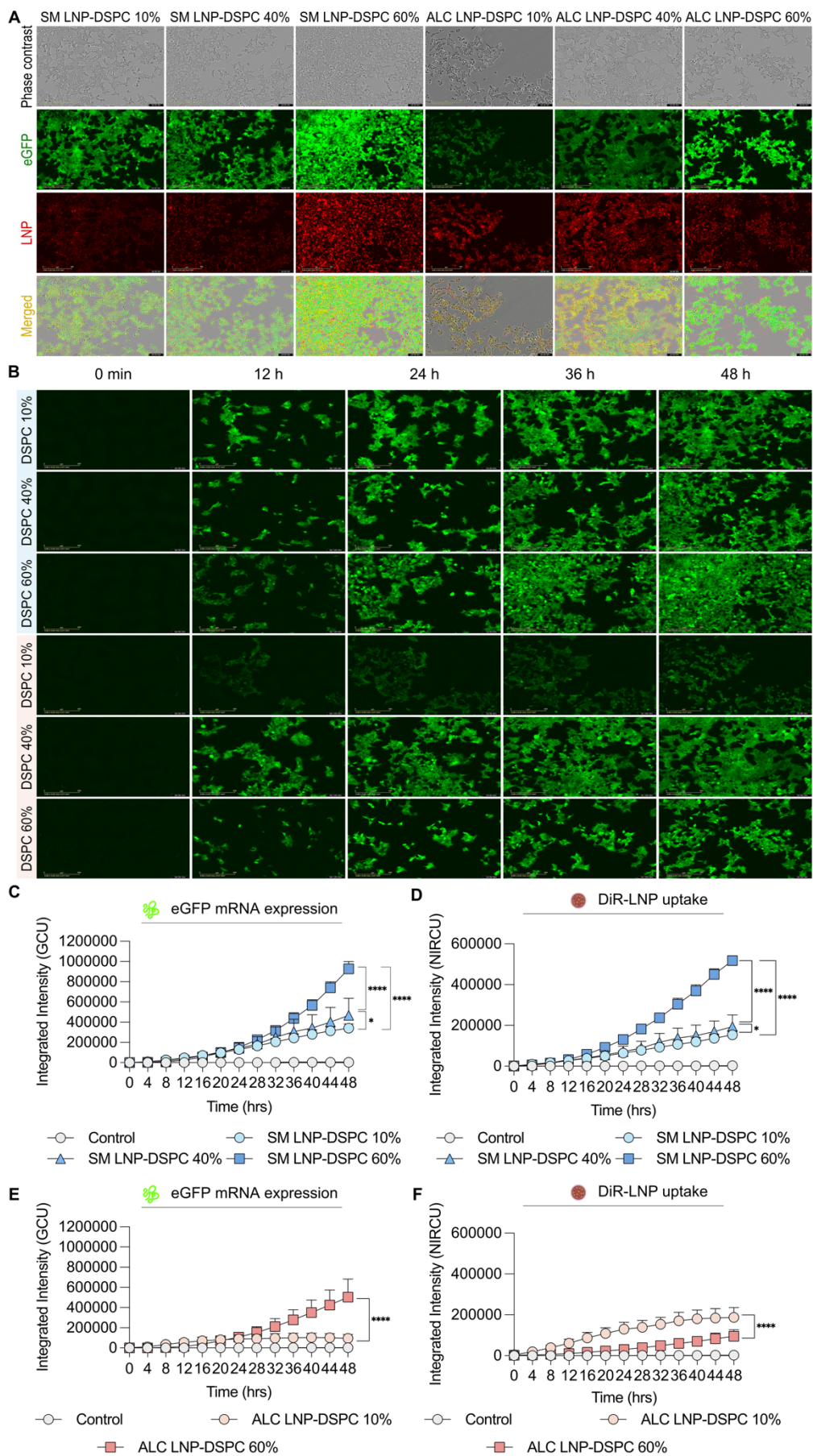

**Figure S8. Enhanced uptake and sustained eGFP expression of non-spherical LNP are maintained across SM- and ALC-LNP.** (A) Representative fluorescence microscopy images showing an increase in LNP uptake and eGFP expression in HEK cells treated with LNP with increasing DSPC content over 48 h of incubation. Scale bar: 200 μm. (B) Representative fluorescence microscopy images of eGFP mRNA expression acquired at 0, 12, 24, 32, and 48 h. (C-F) Quantification of integrated fluorescence intensity over 48 h at 4 h intervals, showing increased eGFP expression and LNP uptake with increasing DSPC percentage. (C,E) Time-lapse quantification of eGFP expression, measured as green channel units (GCU) for LNP formulated with (C) SM and (D) ALC ionizable lipid. (D,F) Time-lapse quantification of DiR-labelled LNP, measured as near-infrared channel units (NIRCU) for LNP formulated with (E) SM and (F) ALC ionizable lipid.

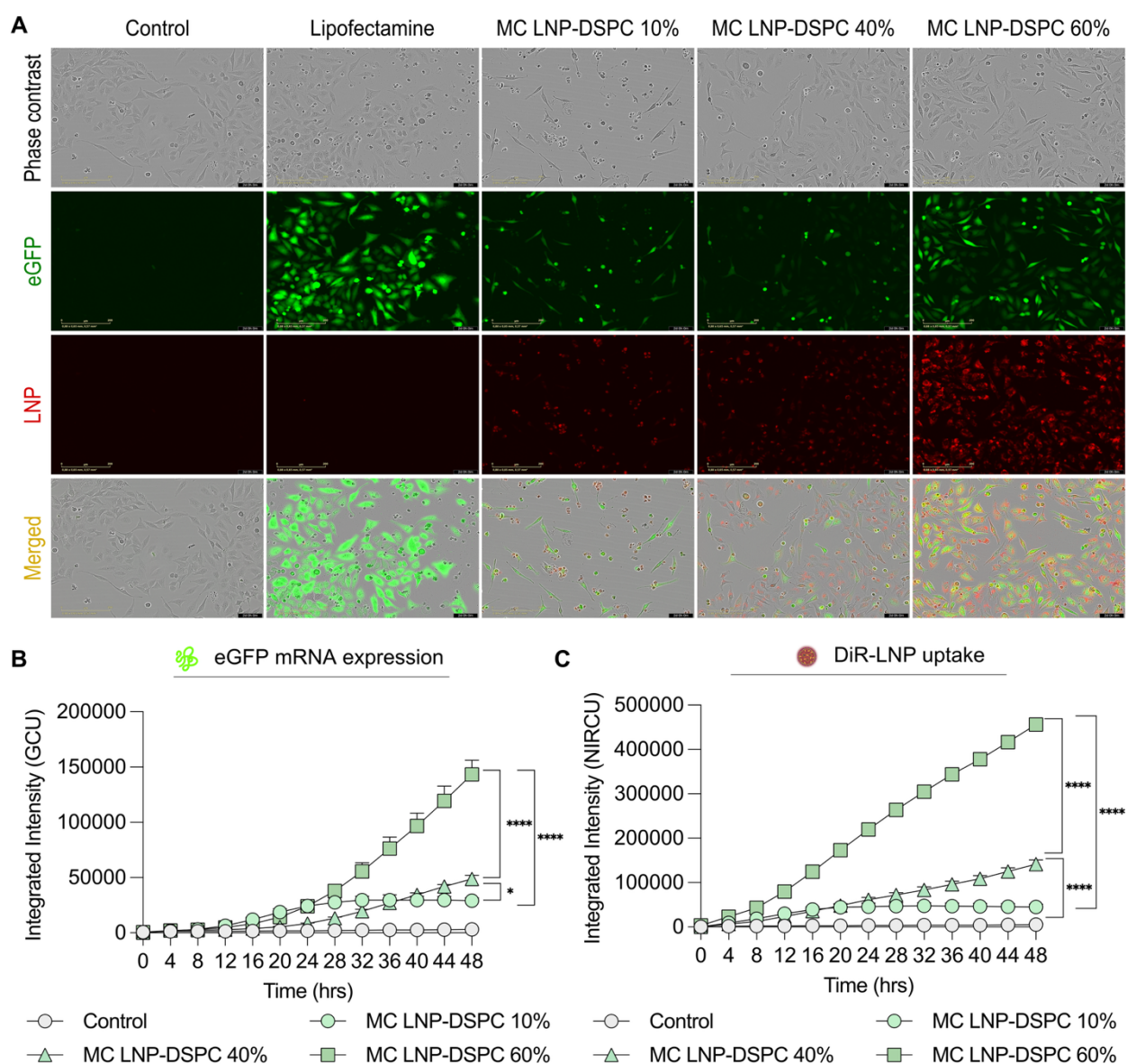

**Figure S9. Non-spherical LNP retain their enhanced uptake and expression profile in HeLa cells.** (A) Representative fluorescence microscopy images of HeLa cells treated with LNP made with MC ionizable lipid and different DSPC levels, captured over 48 hours. Scale bar, 200 μm. (B) Changes in eGFP expression over 48 h, quantified from green channel fluorescence (GCU). (C) Cellular accumulation of DiR-labeled LNP over the same period, quantified from near-infrared fluorescence (NIRCU).

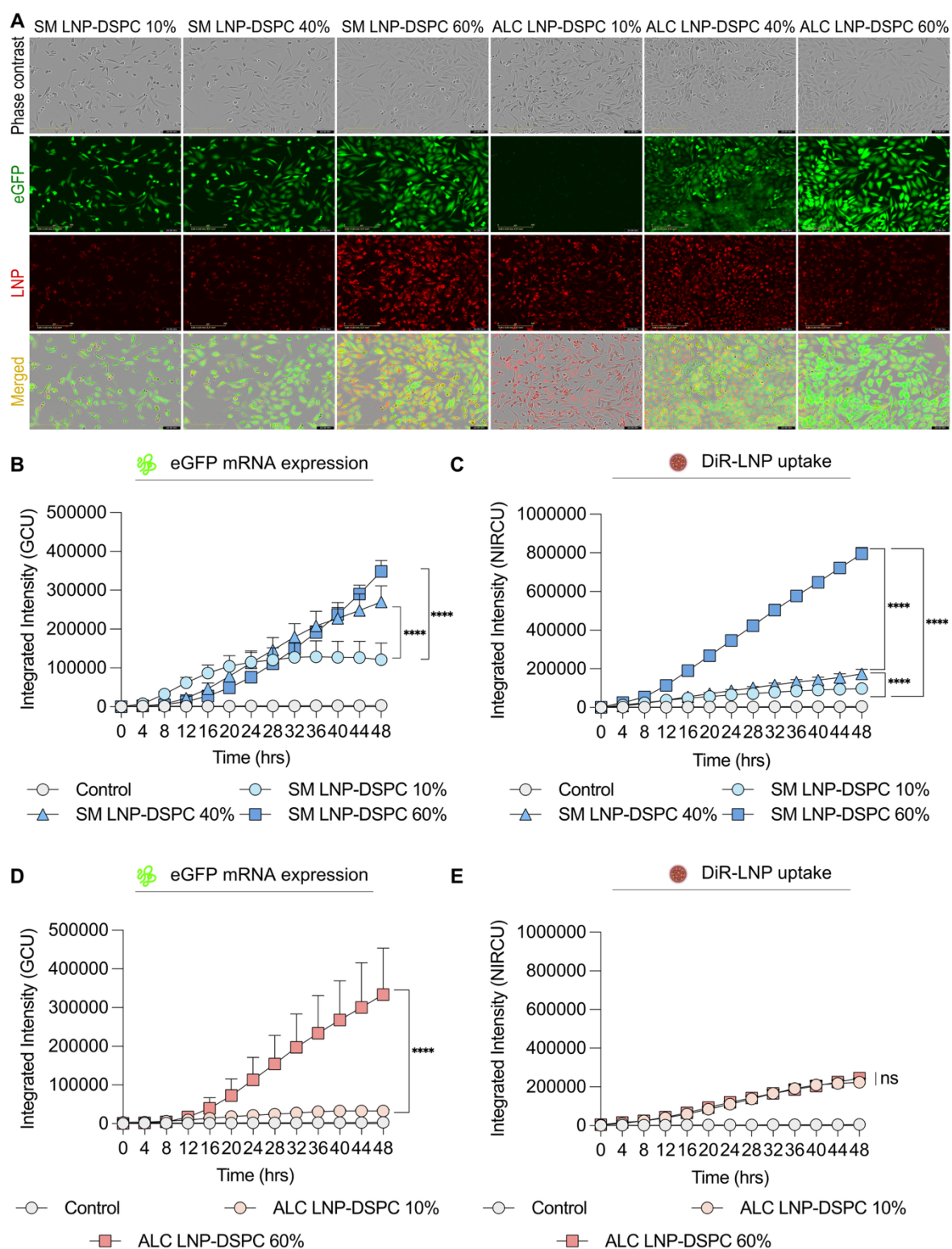

**Figure S10. Sustained eGFP expression by non-spherical LNP persists across ionizable lipid chemistries in HeLa cells.** (A) Representative fluorescence microscopy images showing LNP uptake and eGFP expression in HeLa cells treated with LNP formulated with SM and ALC ionizable lipid and varying DSPC content over 48 h of incubation. Scale bar, 200 μm. (B-E) Quantification of integrated intensity over 48 h, measured at 4 h intervals, showing increased eGFP expression and LNP uptake as DSPC percentage increases. (B,D) Quantification of green channel units (GCU), representing eGFP mRNA expression, for LNP formulated with (B) SM and (D) ALC ionizable lipid. (C,E) Quantification of near-infrared channel units (NIRCU), corresponding to DiR-labeled LNP uptake, for LNP formulated with (C) SM and (E) ALC ionizable lipid.
